# Correcting spatial transcriptomics data affected by a prevalent transcript leakage problem across platforms, species, and tissues

**DOI:** 10.64898/2026.06.13.732076

**Authors:** Christina Huan Shi, Yibo Zhai, Savio Ho-Chit Chow, Liangbang Li, Chase M. Carver, Marcos G. Teneche, Jesus Flores, Colin Kern, Peter D. Adams, Bing Ren, Marissa J. Schafer, Quan Zhu, Yingying Wei, Kevin Y. Yip

## Abstract

Spatial transcriptomics has been widely applied to study the spatial distribution of cell types, cell states, and specific gene expression in tissue samples. However, we show that there is a prevalent transcript leakage problem in spatial transcriptomics data, where transcripts expressed by a cell diffuse to its neighborhood and are recurrently detected in the nearby cells. By analyzing published data sets, we show that this problem is general across data produced from different tissues and different species using different imaging-based and sequencing-based spatial transcriptomics platforms. It affects both upstream tasks such as expression quantification as well as downstream tasks such as cell-type annotation and detection of spatially-dependent gene expression. To tackle the transcript leakage problem, we propose a reference-free Bayesian model-based method, DeLeakage, which cleans up the data much more effectively than existing denoising methods. DeLeakage also improves cell-type annotation and avoids false detection of spatially dependent expression.

## Introduction

Spatial transcriptomics (ST) technologies are rapidly advancing our understanding of the organization of complex tissues in both healthy and diseased states^1-6^. However, in several recent benchmarking studies, transcripts were detected in cell types that they are not expected to be expressed^7-10^. Here we show that a major contributor to transcript detection in unexpected cell types is the leaking of transcripts to surrounding cells during the experimental process. Unfortunately, existing deconvolution methods developed for bulk genomic data^11-19^ or ST data^20-23^ cannot tackle the transcript leakage problem because these methods assume that the observed transcriptome profile of a cell/spot/sample is the result of mixing different cells/cell types at certain proportions, and all genes follow the same proportions (Figure 3B). In contrast, as we show below, different genes have very different leakage properties (Figure 3A). Therefore, a computational method for effectively adjusting ST data affected by transcript leakage is still lacking.

By analyzing a variety of ST data produced from various tissues and species using different imaging-based and sequencing-based ST platforms, we show that transcript leakage is prevalent and various standard analyses are affected. To tackle the problem, we propose a reference-free Bayesian hierarchical model-based method, DeLeakage, to adjust gene expression levels affected by transcripts leaked from neighboring cells through a diffusion process. Our model contains different diffusion parameters for different genes. On the theoretical side of this study, we prove that our model is identifiable, which ensures endogenous expression can be distinguished from the effects of transcript leakage and the model parameters can be estimated reliably. In contrast, classic reference-free deconvolution methods usually build upon non-negative matrix factorization, which is known to be non-identifiable^24^, while reference-based deconvolution methods can be inaccurate when the reference does not provide matched cell types or contain strong batch effects. On the practical side of this study, we show that DeLeakage substantially outperforms existing ST denoising methods on both simulated and real ST data.

## Results

### Evidence of transcript leakage in spatial transcriptomics data

To study why some transcripts are detected in unexpected cell types, we analyzed a published ST data set of the mouse brain^25^ produced using multiplexed error-robust fluorescence in situ hybridization (MERFISH)^26^. We found that transcripts of *Slc17a7*, which should be primarily expressed as a marker of excitatory, glutamatergic neurons and rarely expressed in non-neurons^27^, were frequently detected in non-neuronal cells (74%; Figure 1A). The spatial distribution of them closely resembles the distribution of excitatory neurons (Figure 1B), suggesting potential leakage of *Slc17a7* transcripts from excitatory neurons to nearby cells. Consequently, in the hippocampal region, which has a high density of excitatory neurons (Figure 1B), non-neurons with a high detected level of *Slc17a7* form “contour lines” right next to the principal cells (Figure 1B, inner rectangle), which clearly illustrates the potential transcript leakage.

**Figure 1.**
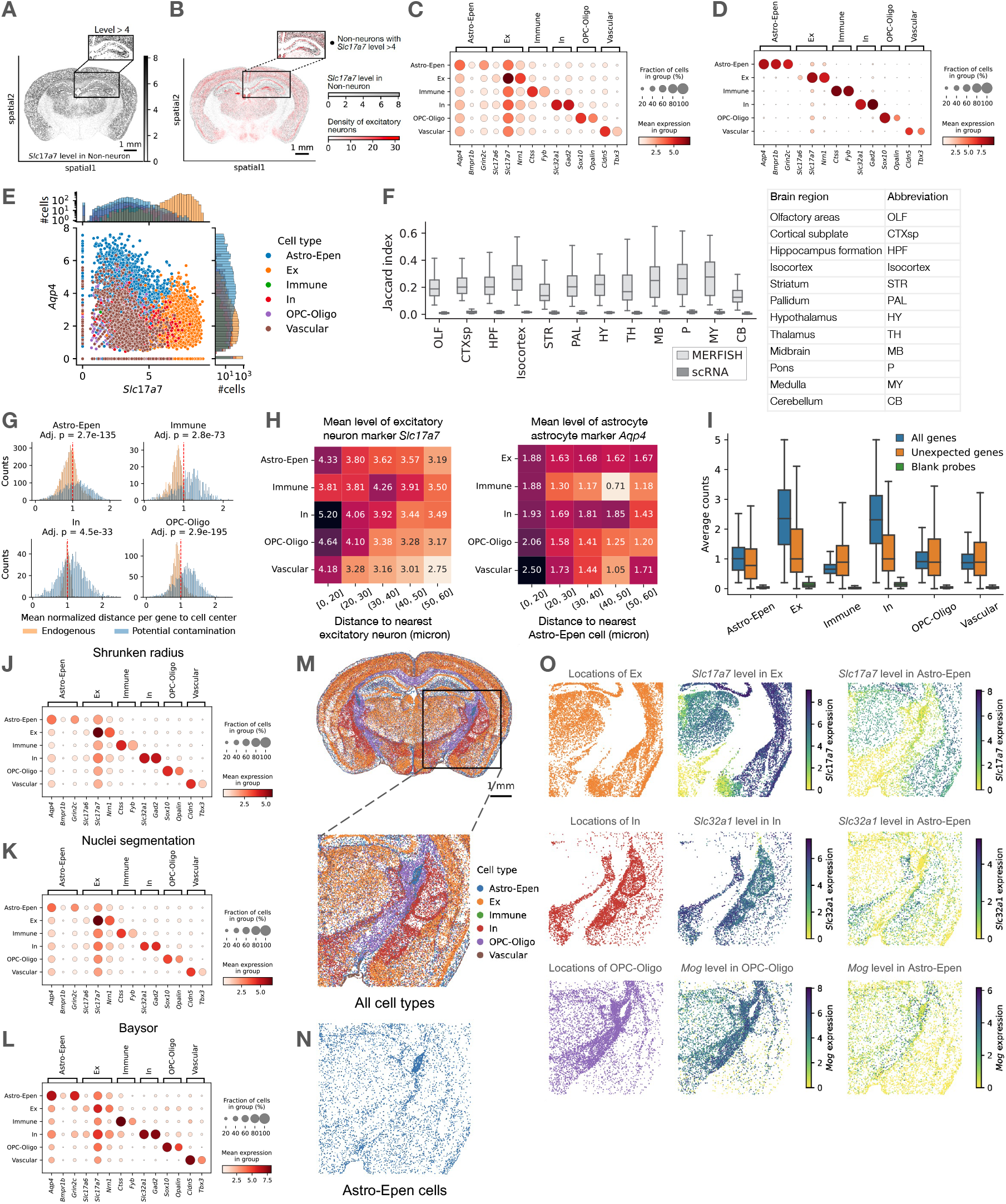
Evidence of transcript leakage in spatial transcriptomics data. (**A**) Spatial distribution of excitatory neuron marker *Slc17a7* transcripts in non-neuronal cells. (**B**) Spatial density distribution of excitatory neurons overlaid on spatial distribution of non-neuronal cells with a high detected level of *Slc17a7*. The inner rectangle shows the hippocampal region. (**C**-**D**) Detection of transcripts of cell type-specific genes in the MERFISH (C) and scRNA-seq (D) data. All results in these and following panels (except Panels F, M, N, and O) were produced using data from the hippocampal formation region. (**E**) Co-detection of excitatory neuron marker *Slc17a7* transcripts and astrocyte marker *Aqp4* transcripts in individual cells in the MERFISH data. (**F**) Co-detection of transcripts in the MERFISH and scRNA-seq data of gene pairs supposed to have mutually exclusive expression in different brain regions. For each gene pair, the Jaccard index is defined as the number of cells with both types of transcripts detected divided by the number of cells with either detected. Each box-and-whisker plot shows the distribution of Jaccard index across all the gene pairs. (**G**) Distribution of endogenous and potential contamination transcripts’ distances from cell center. (**H**) Average detected transcript levels of potential contamination genes in cells at different distances from the nearest potential contamination source. The contamination genes shown are *Slc17a7* with excitatory neurons as potential contamination source (left) and *Aqp4* with astrocytes as potential contamination source (right). (**I**) Average detected transcript counts of all genes, unexpected genes, or negative control blank probes, across all cells. (**J**-**L**) Detection of transcripts of cell type-specific genes in the MERFISH data resulted by three alternative cell segmentation strategies/methods, including shrinking the cell radius by half for the whole cell-body segmentation by CellPose (J), nuclei-only segmentation by CellPose (K) and Baysor segmentation (L). (**M**-**N**) An area for illustrating the effects of transcript leakage on the identification of spatially dependent gene expression patterns, with all major cell types shown (E) or only Astro-Epen cells (F). (**O**) Cell type markers with significant spatially-dependent expression identified in Astro-Epen cells likely caused by transcript leakage. Each row shows the case of one of the cell type markers, with the first column showing the spatial distribution of cells of the expected cell type, which serve as the potential contamination sources, the second column showing the detected levels of the maker in these cells, and the third column showing the detected level of the maker in the Astro-Epen cells.

Beyond *Slc17a7*, transcripts of various other cell type markers are also commonly detected in unexpected cell types (Figure 1C). This more likely represents a technical problem than a biological phenomenon because in dissociative single-cell RNA sequencing (scRNA-seq) data from the same study, cell-specific marker transcripts are much more exclusively detected in the expected cell types (Figure 1D). The unexpected transcript detection is not due to incorrect clustering of cells of different cell types or incorrect cell-type annotations, because expected and unexpected markers are also co-detected in individual cells (Figure 1E). Based on this observation, we compiled a list of gene pairs expected to have mutually exclusive expression in a cell (Methods) and found that transcripts of these gene pairs are detected in the same cell much more frequently in the MERFISH data than in the scRNA-seq data in all brain regions (Figure 1F, Supplementary Figure 1).

Next, we found that in the MERFISH data, transcripts of potential contamination genes are significantly further away from the cell center than endogenously expressed genes (Methods, Figure 1G), which supports our hypothesis that the former transcripts are from an exogenous source. As another important support for our transcript leakage hypothesis, the level of unexpected transcripts in a cell is higher when it is closer to a cell that endogenously expresses it, which acts as the potential contamination source (Figure 1H). This observation also shows that the unexpected transcripts are less likely explainable by contamination from cells at the same 2-dimensional location but residing in other imaging layers (the “z-planes”). We further found that the detected transcript levels of the unexpected genes are much higher than the detected levels of transcripts binding the negative control “blank” probes (Figure 1I), indicating that the unexpected transcript detection cannot be explained by false positive signals or off-target probe binding.

Finally, to test whether the unexpected transcripts are due to cell segmentation errors, we used three other approaches to perform cell segmentation (Methods). In all three cases, transcripts of cell type markers are still observed in unexpected cell types (Figure 1J-L), showing that this issue cannot be fully explained by incorrect cell segmentation.

Together, these results support that transcript leakage is the most plausible explanation for the detection of the unexpected transcripts as compared to the alternative explanations discussed.

Next, we investigated the effects of transcript leakage on the detection of genes with spatially-dependent expression patterns. In an area that consists of multiple regions within the parietal and temporal lobes, including hypothalamus, thalamus, cerebral cortex, and hippocampal formation regions (Figure 1M), the spatial locations of Astro-Epen cells follow a very specific pattern (Figure 1N). We identified genes that show significantly different levels in Astro-Epen cells at different spatial locations.

These genes include markers of some other cell types, including the excitatory neurons marker *Slc17a7*, inhibitory, GABAergic neurons marker *Slc32a1*, and OPC-Oligo marker *Mog*. The Astro-Epen cells with high detected levels of these genes strongly co-localize with cells from these other cell types (Figure 1O), in a way highly independent of the intrinsic spatial distribution of Astro-Epen cells (Figure 1N), indicating that the spatially-dependent levels of these transcripts are more likely caused by transcript leakage than genuine biology.

In the same way, we analyzed several additional ST data sets, including 1) MERFISH data produced from the human heart^28^, 2) Xenium data produced from the mouse brain, and 3) Pixel-seq data produced from the mouse brain^29^. In all three cases, we found the same evidence of transcript leakage, with a Jaccard Index of co-detection of mutually exclusive gene pairs from 0.195 to 0.353 (Figure 2, Supplementary Results), showing that it is a prevalent problem in ST data.

**Figure 2.**
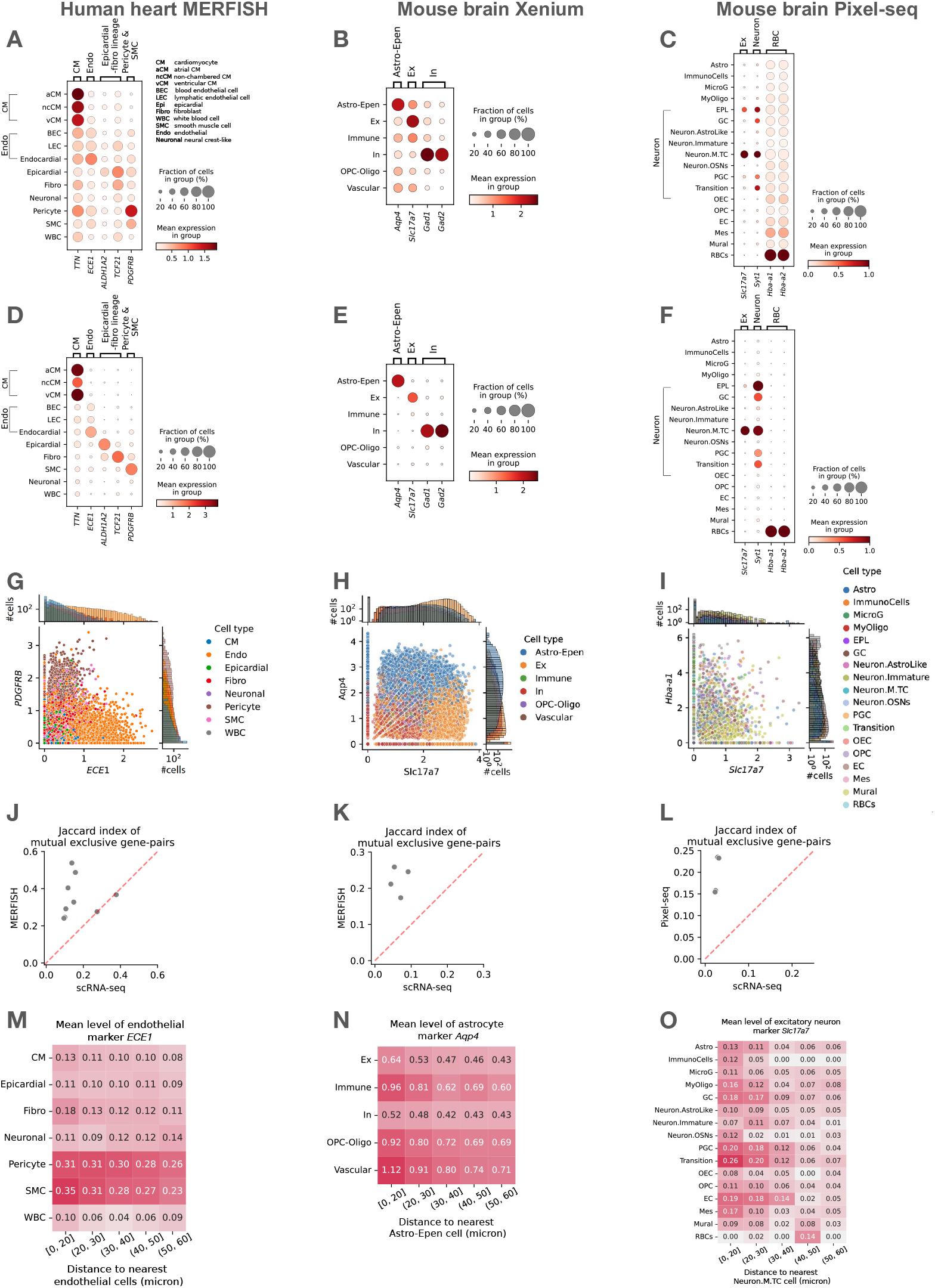
Generality of transcript leakage across data produced from different tissues of different species on different spatial transcriptomics platforms. (**A-C**) Detection of transcripts of cell type-specific genes in the spatial transcriptomics data. In the human heart MERFISH data set (A), aCM, ncCM, and vCM are all subtypes of cardiomyocytes, while BEC, Endocardial, and LEC are all subtypes of endothelial cells. These subtypes share some common markers. Similarly, in the mouse brain Pixel-seq data set, some neuron subtypes share some common markers. (**D-F**) Detection of transcripts of cell type-specific genes in the corresponding dissociative scRNA-seq/snRNA-seq data. (**G-I**) Co-detection of markers of two different cell types in individual cells in the spatial transcriptomics data. (**J-L**) Co-detection of transcripts in the spatial transcriptomics and dissociative scRNA-seq/snRNA-seq data of gene pairs supposed to have mutually exclusive expression. For each gene pair, the Jaccard index is defined as the number of cells with both types of transcripts detected divided by the number of cells with either detected. (**M-O**) Average detected transcript level of a potentially contamination gene in cells at different distance from the nearest potential contamination source.

### DeLeakage: A statistical method for correcting for transcript leakage

To tackle the transcript leakage problem, we propose a reference-free Bayesian hierarchical model-based method, DeLeakage (Decontamination for Transcript Leakage), which decomposes the observed transcript counts of each cell into its endogenous expression and transcripts leaked from other cells through a diffusion process (Figure 3A). Our model includes a different diffusion parameter for each gene, thus allowing the capture of gene-specific properties that affect leakage patterns. This is different from the conventional deconvolution approaches, which model the observed expression profile of a sample as the result of mixing different cell types without considering gene-specific parameters (Figure 3B).

**Figure 3.**
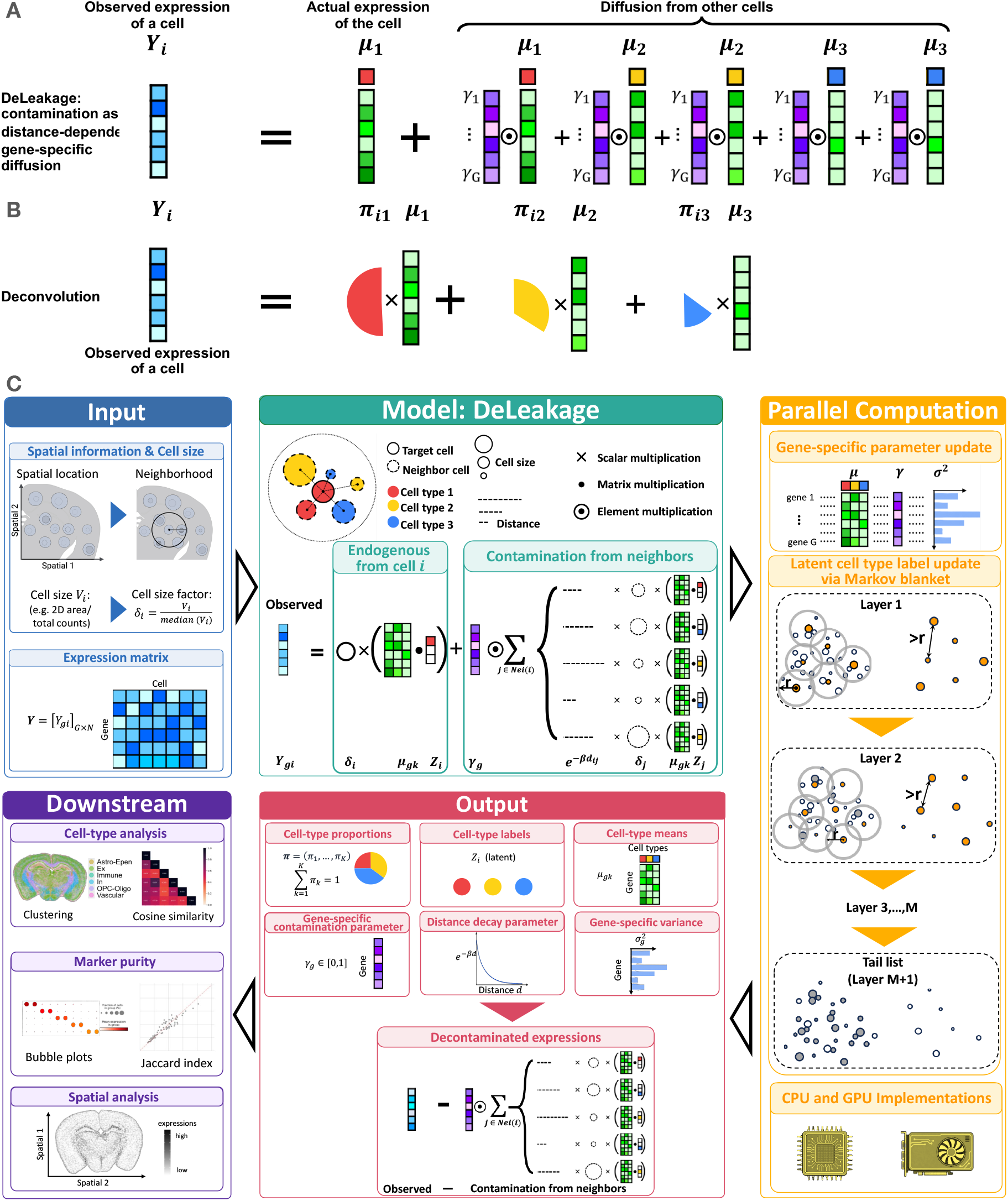
The DeLeakage method. (**A**) The DeLeakage approach, in which the observed expression level of a gene in a cell is modeled as the sum of its actual expression and expression of the gene in neighboring cell diffused to the cell according to their distance and gene-specific properties. (**B**) The conventional deconvolution-based denoising approach, in which the observed expression profile of a cell is modeled as the mixture of expression profiles of different cell types. (**C**) Illustration of the DeLeakage method. The input includes the spatial location, estimated size, and observed expression profile of each cell. The DeLeakage model contains parameters related to size of each cell *δ*_*i*_, mean expression of each gene in each cell type *μ*_*gk*_, latent cell type of each cell *Zi*, proportion of cells belonging to each cell type *π*_*k*_, diffusion coefficient of each gene *γ*_*g*_, and distance-dependent decay *β*. These parameters are estimated from the data using a parallel algorithm for both the CPU and GPU versions. The output of DeLeakage includes the estimated values of the model parameters as well as decontaminated expression profile of each cell. These results can be used for various downstream analyses.

DeLeakage works as follows (Figure 3C, Methods). Suppose that after cell segmentation, we obtain a total of *N* cells and cell *i* has a volume of *V*_*i*_. Denote the number of transcripts of gene *g* assigned to cell *i* as *Y*_*gi*_ . Based on the spatial location of each cell *i*, the subset of other cells in its spatial neighborhood is denoted as *Nei*(*i*). The distance between cell *j* and cell *i* is denoted as *d*_*ij*_. Assume there are *K* cell types in total and the probability for cell *i* to belong to cell type *k* is *Pr*(*Z*_*i*_ = *k*) = *π*_*k*_, where *Z*_*i*_ is the latent cell-type label for cell *i*, and any two cells’ cell-type labels are independent. For cell *i*, given its cell-type label *Z*_*i*_ and the cell-type labels of its neighbors, *Z*_*Nei*(*i*)_, we assume that *Y*_*gi*_ follows a Gaussian distribution 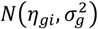 with a mean *η*_*gi*_ and a gene-specific variance 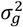. The mean level can be further decomposed into 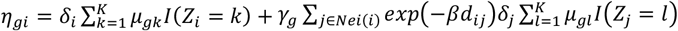, where *μ*_*gk*_ is the mean expression level of gene *g* in cell type *k, γ*_*g*_ ∈ (0,1) is a gene-specific contamination parameter, *β* > 0 is a parameter that specifies the decay of contamination with respect to distance, and 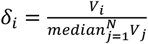 accounts for the impact of cell volume, namely larger cells are likely to leak more transcripts to neighboring cells. Consequently, 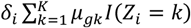 represents cell *i*’s endogenous expression level of gene *g* after accounting for its cell volume, whereas for a neighboring cell *j* ∈ *Nei*(*i*) that belongs to cell type 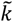, its contribution to cell *i*’s detected transcript level of gene *g* due to transcript leakage is 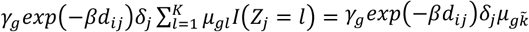, which depends on its cell type 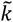, cell volume factor *δ*_*j*_, distance *d*_*ij*_ to cell *i*, and gene *g*’s contamination parameter *γ*_*g*_ .

Theorem 1 below states that DeLeakage’s model is identifiable up to label switching of cell types, thanks to (a) the cellular resolution and (b) spatial structure of ST data under conditions that are very easily met in reality.

#### Theorem 1.

*Given that*

*(C1) For each cell type k, there exists a gene g*^(*k*)^ *such that* 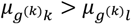 *for all l* ≠ *k; and*

*(C2) There exist three cells i*_1,_ *i*_2,_ *i*_3_ *such that the function* 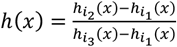 *is strictly monotonic in x, where* 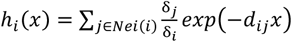 *represents the spatial density of neighbors for cell i, which is normalized by each neighbor’s cell volume and weighted by each neighbor’s distance to cell i with x denoting the scale parameter of the exponential kernel for weighting, then the DeLeakage model is identifiable (up to label switching) in the sense that if Pr*(***Y***|***Θ***) = *Pr*(***Y***|***Θ***^∗^) *for any* ***Y*** *for two sets of parameters* ***Θ*** = {π, (***μ***_1_, …, ***μ***_*K*_), ***γ***, *β*, ***∑***} *and* 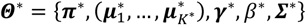, *then* 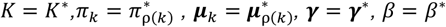, *and* ***∑*** = ***∑***^∗^, *where ρ is a permutation of* {1, …, *K*}.

Condition (C1) requires each cell type to have at least one marker gene that is expressed at a higher level in that cell type than in any other cell type, which is always true in practice. In fact, researchers very often deliberately include marker into the gene panel for imaging-based ST. Condition (C2) is an easily met condition on spatial heterogeneity.

Based on the estimated parameters 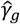 and 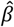, and *μ*_*gk*_ and the inferred cell type labels 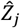, we compute corrected transcript level for gene *g* in cell *i* as 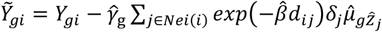 in the same spirit as the batch effect adjustment done in previous studies^30, 31^.

We implemented a CPU version and a GPU version of our Markov chain Monte Carlo (MCMC) algorithm, both conducting parallel computing by partitioning cells into layers so that cells of the same layer are not within each other’s Markov blanket and hence their cell-type labels can be updated in parallel to speed up the calculations (Figure 3C; Methods).

### Validation and benchmarking of DeLeakage using simulated data

To test DeLeakage, we first produced a simulated data set that mimics real ST data (Figure 4A-B, Methods). When we applied DeLeakage to the data, it successfully identified the real cell types (Figure 4C), estimated the real mean expression level of each gene in each cell type (Figure 4D), and estimated the gene-specific diffusion parameters accurately (Figure 4E).

**Figure 4.**
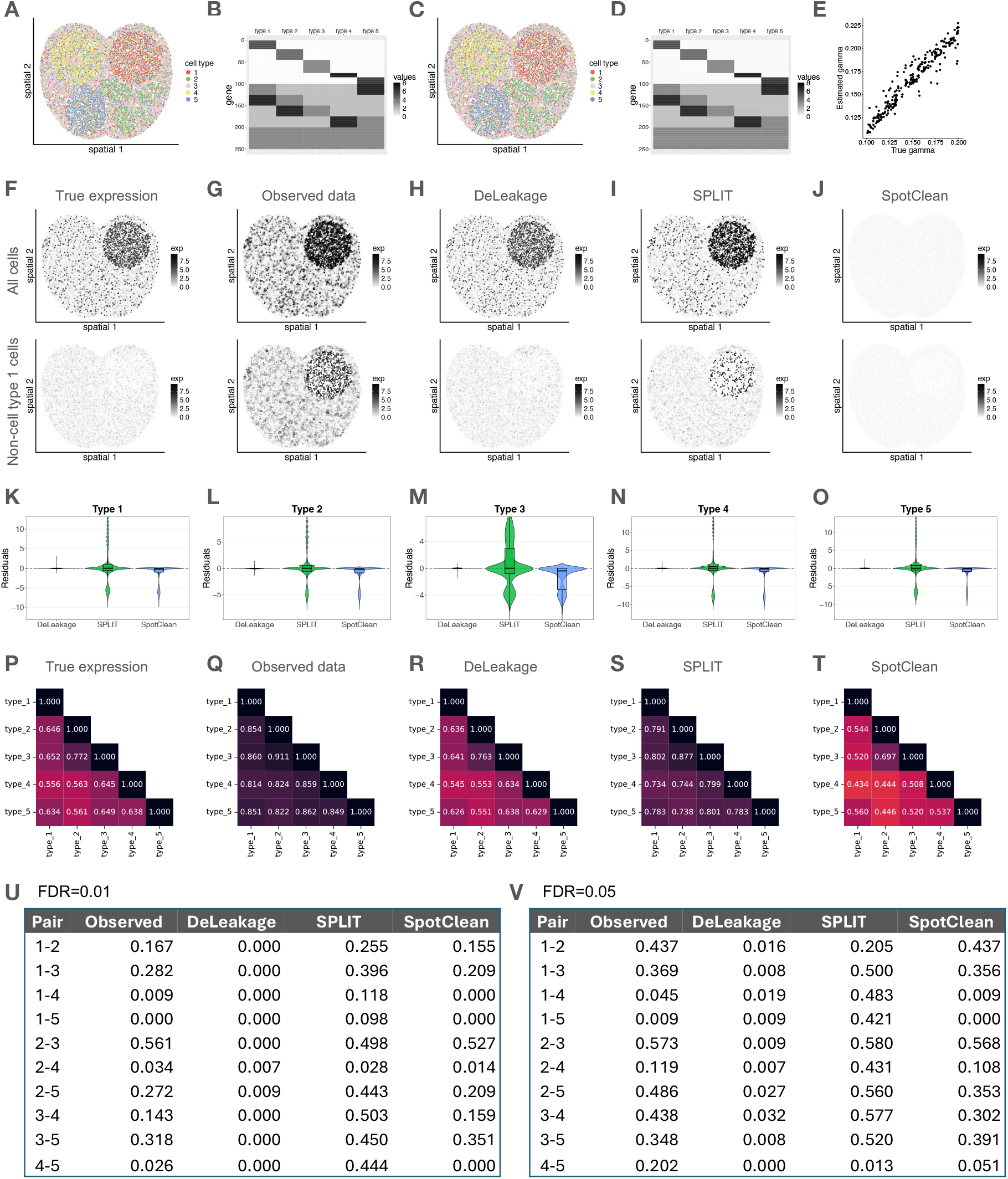
Validation and benchmarking of DeLeakage using simulated data. (**A**) Actual locations of cells in the 5 cell types. (**B**) Actual mean expression level of each gene in each cell type. (**C**) Cell type of each cell inferred by DeLeakage based on the learned values of the latent cell type variables. (**D**) Inferred mean expression level of each gene in each cell type by DeLeakage. (**E**) Comparing DeLeakage-inferred and actual contamination parameter values. (**F**-**J**) Level of one of the marker genes in cell type 1 selected randomly in the initial simulated data before transcript leakage (F), final simulated data after transcript leakage (G), the adjusted data after applying DeLeakage (H), SPLIT (I), or SpotClean (J). In each panel, the upper plot includes all cells, and the lower plot includes only cells not of cell type 1. (**K**-**O**) Residuals between inferred and actual average expression levels of the randomly selected marker gene of each cell type, with the inference performed by DeLeakage, SPLIT, or SpotClean. Each panel shows the result of one cell type. (**P**-**T**) Average cosine similarity between cells from the same cell type or different cell types, based on the actual gene expression levels (U), observed levels affected by transcript leakage (V), and inferred expression levels by DeLeakage (W), SPLIT (X), or SpotClean (Y). (**U**-**V**) False discovery rates of differentially expressed genes between two cell types based on the observed transcript levels or the inferred expression levels by DeLeakage, SPLIT, or SpotClean. The two panels show results when the Benjamini-Hochberg procedure was used to control the false discovery rate at 0.01 (U) or 0.05 (V).

In the data, marker gene of cell type 1 have much higher actual expression levels in these cells (Figure 4F upper) than in other cells (Figure 4F lower), but due to transcript leakage, in the observed data, some cells not of cell type 1 but residing in its spatial territory also have high detected levels of the cell type marker (Figure 4G). After applying DeLeakage, the contamination was successfully removed while retaining true expression levels of the cell type marker (Figure 4H). These findings were also obtained from the other cell types (Supplementary Figures 3-4).

Although no previous methods were specifically designed to tackle transcript leakage, we tested whether the effects of transcript leakage can also be removed using a general ST data denoising method. Therefore, we applied SPLIT^23^ and SpotClean^20^ to our simulated data (Methods). SPLIT first runs RCTD^21^ with reference scRNA-seq data to deconvolute the observed expression levels and estimate the proportions of the primary and the secondary cell types. Subsequently, SPLIT scales the observed expression to a purified expression level with the scaling factor being the primary cell type’s expected fraction of transcripts. SpotClean is a method that leverages background spots that are supposed to have no expression to remove contamination due to messenger RNAs bleed for spot-based ST methods. When we applied these methods to our simulated data, SPLIT failed to remove most contamination (Figure 4I), while SpotClean inferred expression levels much lower than the real levels (Figure 4J). These findings were also obtained from the other cell types (Supplementary Figures 3-4). When we computed the residuals between the inferred and actual gene expression levels of the marker genes (Figure 4K-O), in all five cell types DeLeakage had a residual close to 0 for most genes, confirming the identifiability of our model and the accurate estimation of model parameters by our algorithm. In comparison, in each cell type, SPLIT led to some cells having a large positive or negative residual, while SpotClean over-corrected the data and consistently produced negative residuals.

We also found that DeLeakage was more effective than SPLIT and SpotClean in recovering actual differences between different cell types (Figure 4P-S) and avoiding false positives when calling differentially expressed genes between cell clusters (Figure 4U-V) (Supplementary Results).

Together, these results validate DeLeakage’s model and algorithm and also confirm the necessity for a new method specifically designed for tackling the transcript leakage problem.

### Effective correction of real spatial transcriptomics data using DeLeakage

Next, we applied DeLeakage to the real ST data sets. In the scRNA-seq data of mouse brain, each cell type contains marker genes that are almost exclusively expressed in this cell type (Figure 5A). In contrast, in the original uncorrected MERFISH data, transcripts of various cell type markers are commonly detected in other cell types (Figure 5B). After applying DeLeakage, these cell type markers became more specifically detected in the expected cell types (Figure 5C). In comparison, after applying SPLIT, non-specific detection of cell type markers was still quite prevalent (Figure 5D). For example, among cells other than excitatory neurons, *Slc17a7* was detected in 55% of them in the raw MERFISH data, 21% after applying DeLeakage, and 55% after applying SPLIT.

**Figure 5.**
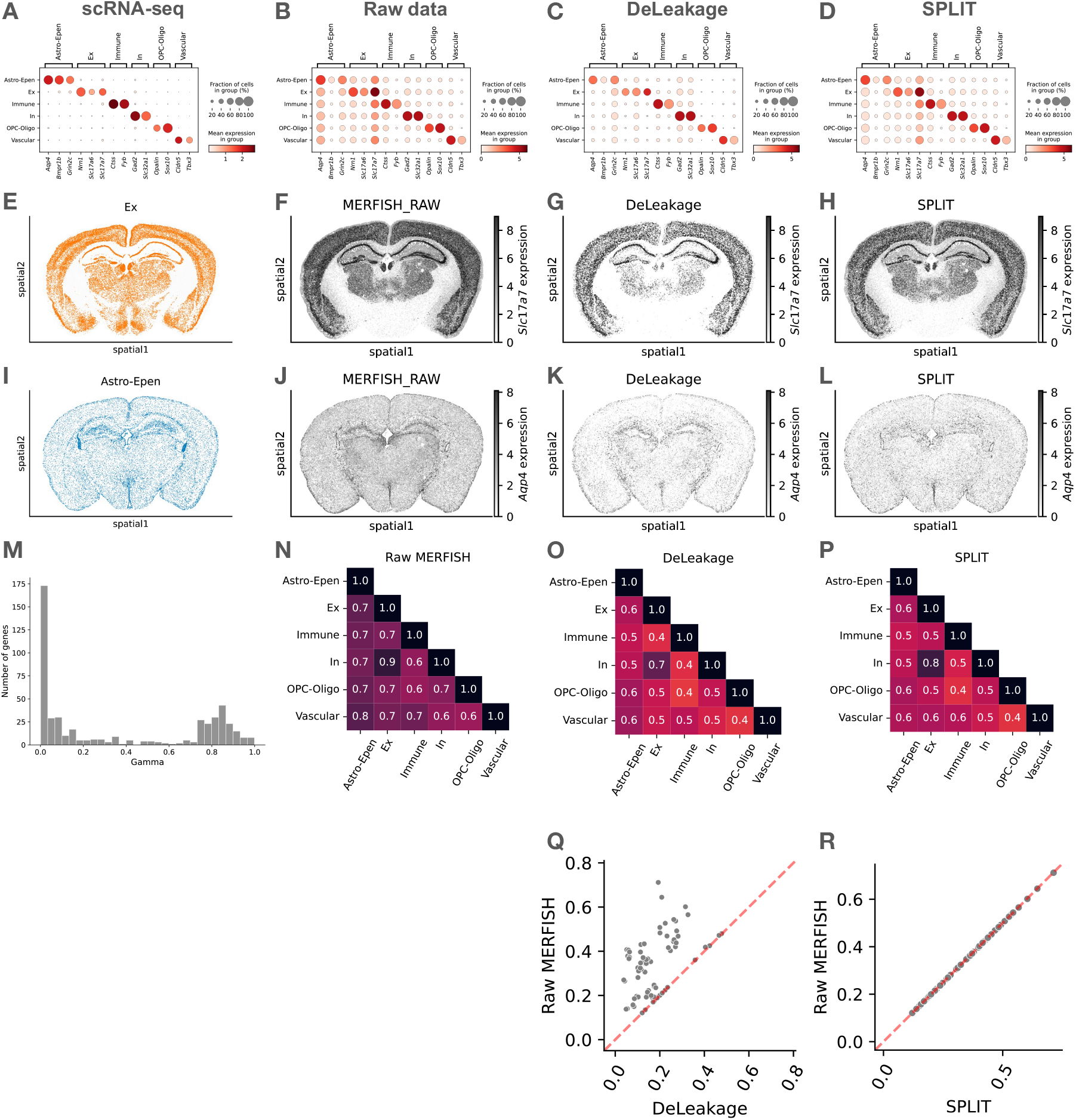
Improvement of transcript quantification in the mouse brain MERFISH data achieved by DeLeakage. (**A**-**D**) Detection of transcripts of cell type-specific genes in the scRNA-seq data (A), raw MERFISH data (B), or after applying DeLeakage (C) or SPLIT (D). (**E-H**) Spatial distribution of excitatory neurons (E) and transcripts of its marker *Slc17a7* in the raw data (F), or after applying DeLeakage (G) or SPLIT (H). (**I-L**) Spatial distribution of Astro-Epen cells (I) and transcripts of its marker *Aqp4* in the raw data (J), or after applying DeLeakage (K) or SPLIT (L). (**M**) Distribution of all genes’ diffusion parameters identified by DeLeakage. (**N**-**P**) Cosine similarity of average gene expression among cell types, based on the raw data (N), or after applying DeLeakage (O), or SPLIT (P). (**Q-R**) Co-detection of transcripts of gene pairs supposed to have mutually exclusive expression before and after applying DeLeakage (Q), or SPLIT (R). For each gene pair, the Jaccard index is defined as the number of cells with both types of transcripts detected divided by the number of cells with either detected.

As illustrative visual examples, in the raw MERFISH data, the excitatory neuron marker *Slc17a7* was detected in regions with a low density of excitatory neurons (Figure 5E,F). After applying DeLeakage, *Slc17a7* expression was detected primarily in excitatory neurons (Figure 5G). Of note, DeLeakage did not take cell-type annotations or information about marker genes as its input. In contrast, after applying SPLIT, detection of *Slc17a7* in these other cell types was still prevalent (Figure 5H). Similarly, after applying DeLeakage, detection of *Aqp4* became highly consistent with locations of Astro-Epen cells (Figure 5I-L).

In these visualizations, *Slc17a7* appears to diffuse further than *Aqp4*. To see whether DeLeakage was able to capture the different levels of diffusion of different genes’ transcripts, we examined the diffusion parameters learned by the model. We found that the diffusion parameters took on a wide range of values, with a mean of 0.37 and a standard deviation of 0.38 (Figure 5M), confirming that it is important to include these gene-specific diffusion parameters in the DeLeakage model.

In the raw data, due to transcript leakage, cells from different cell types were fairly similar (Figure 5N). After applying DeLeakage, the different cell types became more distinguishable (Figure 5O). SPLIT was also able to achieve this but to a slightly lower extent (Figure 5P).

Finally, we used the scRNA-seq data to define gene pairs expected to have mutually exclusive expression (Methods). After applying DeLeakage, co-detection of these gene pairs in the same cell was reduced (Figure 5Q). In contrast, SPLIT did not reduce co-detection of these gene pairs because it maintained a positive expression value of all the originally detected genes (Figure 5R).

SpotClean was excluded from the above comparisons because it was too computationally demanding to be applied to the whole mouse section. Therefore, we cropped a small area and applied DeLeakage, SPLIT, and SpotClean to only cells within this region (Methods, Supplementary Figure 5A). DeLeakage again achieved better data adjustments as evidenced by more specific marker expression (Supplementary Figure 5B-O), better separation of cell types (Supplementary Figure 5P-S), and lower co-detection of mutually exclusive transcripts (Supplementary Figure 5T-V).

We also applied DeLeakage to the other three ST data sets and found that it successfully adjusted the data (Supplementary Figures 6-8, Supplementary Results).

### Improvement of downstream analyses due to data adjustments by DeLeakage

Next, we investigated whether DeLeakage could improve cell-type annotation of cell clusters (Methods). The results show that in all 4 data sets, the cell clusters produced from DeLeakage-adjusted data were more consistent with the cell-type annotations than clusters produced from the raw data, with an increase of Adjusted Rand Index (ARI) by 71.7% (Figure 6A). In comparison, SPLIT-adjusted data only led to small improvements, with an increase of ARI by 14.2% (Figure 6A). Correspondingly, locations of cells in different cell types also became more consistent with the annotations after applying DeLeakage (Figure 6B). Since the cell-type annotations were produced using both ST and dissociative scRNA-seq data together (Methods), the improvement achieved by DeLeakage has made cell-type annotation based on ST data alone easier.

**Figure 6.**
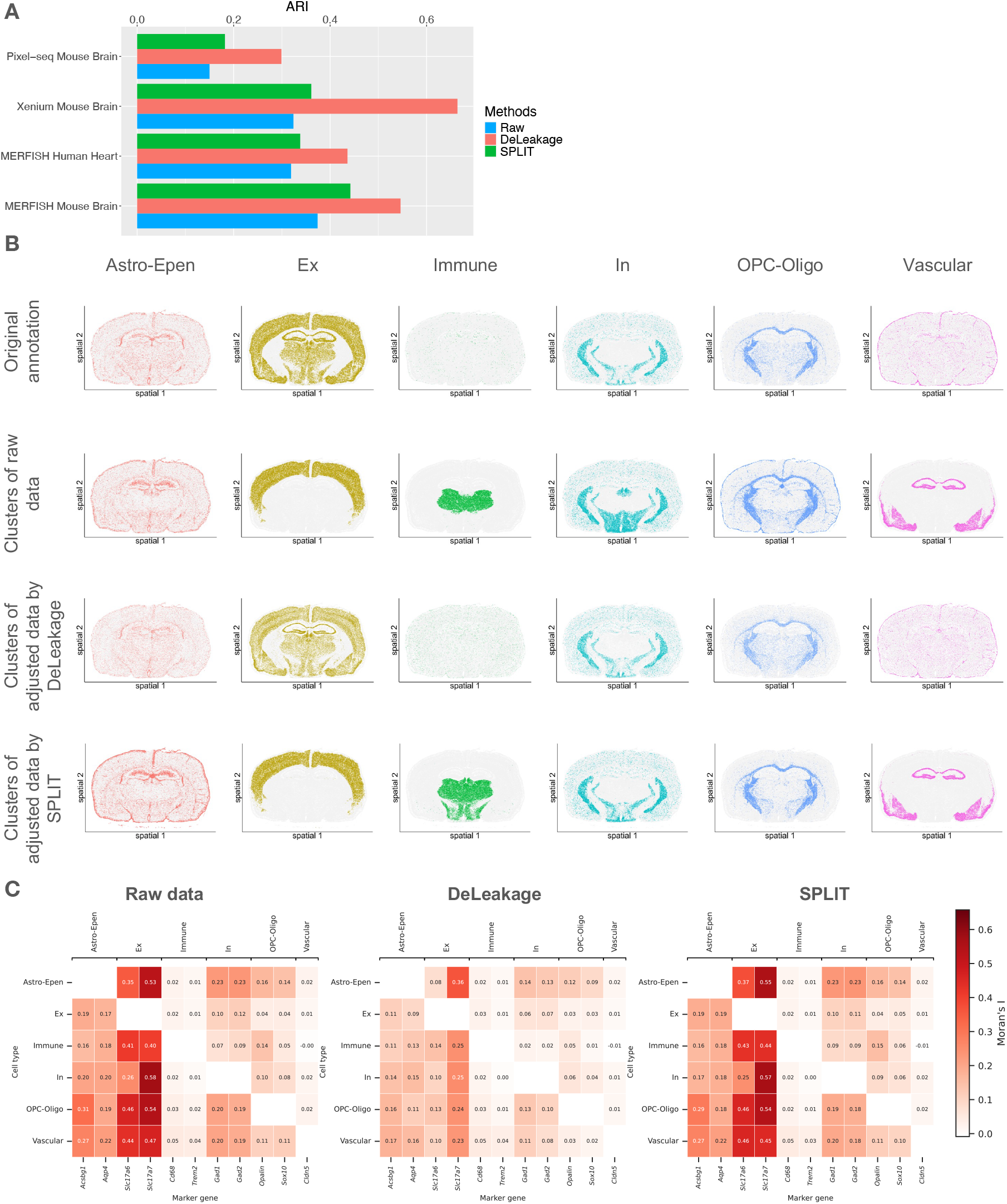
Improvement of downstream analyses achieved by DeLeakage. (**A**) Consistency between cell clusters and cell-type annotations, based on clusters produced from raw data, DeLeakage-adjusted data, or SPLIT-adjusted data. (**B**) Locations of cells in different cell types (columns) in the mouse brain Xenium data, based on the cell-type annotation (first row), clustering of raw data (second row), clustering of DeLeakage-adjusted data (third row), or SPLIT-adjusted data (fourth row). (**C**) Spatial auto-correlation of unexpected transcripts in the mouse brain Xenium data, based raw data (first column), DeLeakage-adjusted data (second column), or SPLIT-adjusted data (third column).

We also quantified spatially dependent expression in unexpected cell types by spatial auto-correlation (Methods). When a gene is not expected to be expressed in a cell type, the detected transcript levels in the cells of this cell type should reflect mostly noise and therefore should not show strong spatial auto-correlation. Indeed, after applying DeLeakage, such spatial auto-correlation was reduced (Figure 6C, Supplementary Figure 9), but SPLIT did not consistently achieve that (Figure 6C).

Finally, we evaluated the running time and peak RAM usage of DeLeakage and found it to be highly scalable (Supplementary Figure 10, Supplementary Results).

## Discussion

To the best of our knowledge, DeLeakage is the first method that can estimate the true underlying expression levels and the levels of contamination due to transcript leakage for ST data. DeLeakage allows the contamination level of transcript leakage to be gene-specific, which is infeasible by common deconvolution methods. As DeLeakage does not require any reference data as input, it avoids problems such as batch effects and missing cell types suffered by reference-based deconvolution methods. On the theoretical side, we prove that DeLeakage is identifiable because of the spatial heterogeneity of neighboring patterns. Computationally, our GPU-parallelized MCMC algorithm enables very efficient computation for the large data volume of spatial transcriptomics data. Empirically, applying to both simulated and real data sets, we showed that DeLeakage can effectively remove effects of transcript leakage, leading to more accurate quantification of expression levels and less bias in various downstream analyses. Due to the lack of gene-specific diffusion parameters and/or reliance on an external reference, existing methods were not able to clean up the transcript leakage effects as effectively.

In this study, we use the Gaussian distribution to model the conditional distribution of the observed expression data, *Y*_*gi*_ |*Z*_*i*_, ***Z***_*Nei*(*i*)_, because of its computation convenience and reasonable decontamination performance. For a more delicate modeling, we can replace the Gaussian distribution with the negative binomial or the zero-inflated negative binomial distribution to further model the count data nature of spatial transcriptomics data, though at the sacrifice of computation efficiency. In addition, the Potts model can be incorporated to model any spatial dependencies of the cell types.

The current DeLeakage model uses cells as basic units, where the contamination transcripts received by a cell are leaked by other cells nearby. As a result, transcripts not assigned to any cell are ignored. These transcripts could potentially be also informative. For example, a lysed cell can easily leak out transcripts to the surrounding, but the cell segmentation algorithm may fail to identify it as a cell. Therefore, it would be useful to explore ways to include these transcripts in modeling transcript leakage. This approach may also alleviate errors made by cell segmentation algorithms.

In this study, we focused on tackling the transcript leakage problem and relied on standard methods to handle other ST data processing tasks. If transcript leakage also has a strong effect on these tasks, it could be useful to incorporate DeLeakage as part of the overall data processing model rather than a separate step. Given the prevalence of transcript leakage across tissue types and spatial transcriptomic platforms, we envision that DeLeakage will be widely adopted in the analysis of spatial transcriptomic data and greatly reduce the risk of data misinterpretation.

## Methods

### Real data sets

#### The mouse brain MERFISH data set

The processed mouse brain MERFISH data^25^ was downloaded following the descriptions and instructions provided at https://github.com/AllenInstitute/abc_atlas_access/blob/main/descriptions/MERFISH-C57BL6J-638850.md. The MERFISH data set was generated using a 500-gene panel, and it included serial coronal slices spanning the entire mouse brain. In the original study, the cell types of the cells in the MERFISH data were annotated mapping each of them to the transcriptomic taxonomy defined by the scRNA-seq data. We used these cell-type annotations directly in our analyses. For the MERFISH data, normalized expression levels were computed by first dividing raw counts in each cell by its cell volume, multiplying the result by 1,000, and then taking log2 after adding one as a pseudo-count.

We used data from slice #38, which was cut from the middle and covers all brain regions clearly. It contained 130,112 cells that passed quality control (QC).

The companion scRNA-seq data from the same study was downloaded following the instructions at https://github.com/AllenInstitute/abc_atlas_access/blob/main/descriptions/WMB-10X.md. Over 700 scRNA-seq libraries were generated from anatomically defined tissue microdissections of different brain regions. In our analysis, we used scRNA-seq data generated by the 10x v3 protocol to minimize potential protocol-induced confounding effects, which contained around 2.3 million cells that passed QC. The raw counts and log-normalized gene expression values provided by the original study were used in analysis and visualization. We used the cell-type annotations of the scRNA-seq data provided by the original study in our analyses.

#### The human heart MERFISH data set

The processed human developing heart MERFISH data and companion scRNA-seq data^28^ were downloaded from https://cells.ucsc.edu/?ds=hoc. The MERFISH data was generated using a 238-gene panel. We analyzed 72,962 cells that passed QC. This study also generated companion scRNA-seq data with 142,946 cells that passed QC. The scRNA-seq data from samples collected at 13 post-conception weeks, similar to the MERFISH samples, were used. In the original study, cell types of the cells in the MERFISH data were annotated by clustering them based on their gene expression profiles followed by assigning cell type labels to the clusters based on marker genes and corresponding scRNA-seq data. We used the cell-type annotations of scRNA-seq and MERFISH data provided by the original study in our analyses. To normalize MERFISH data, the raw counts were divided by cell volume per cell and then scaled up by multiplying 1,000, followed by log2 with one as pseudo-count. For scRNA-seq data, the raw counts and log-normalized gene expression values provided by the original study were used in analysis and visualization.

#### The mouse brain Xenium data set

The mouse brain Xenium data of a coronal slice was downloaded from the 10x Genomics web site https://www.10xgenomics.com/datasets/fresh-frozen-mouse-brain-for-xenium-explorer-demo-1-standard on July 18th, 2025. The downloaded data contain raw cell x gene expression matrix and meta data files of 248 genes and 130,870 cells. We performed stringent QC of this data set as follows. We filtered out low-quality cells with less than 10 genes with non-zero expression or less than 10 read counts, and we also filtered out cells of cell area less than 25 mm^2^ or at least 3 times of the medium cell area. Potential doublets were identified using Scrublet (v0.2.3)^32^ and cells with doublet score at least 0.2 were removed. The remaining 129,788 cells were annotated by scANVI (packed in scvi-tools v1.3.2)^33^ using an annotated mouse brain scRNA-seq data^25^ as reference. The Xenium raw counts were first normalized by the total number of transcripts using the “normalize_total” function, and then log-normalized using “log1p” with “base=2” using the Scanpy package (v 1.10.3)^34^.

#### The mouse brain Pixel-seq data set

The processed mouse brain (olfactory bulb region) Pixel-seq data^29^ was downloaded from Gene Expression Omnibus (accession number: GSE186097). The corresponding cell-type annotation information was downloaded from the GitHub repository at https://github.com/GuLABatUW/Pixel-seq. This ST data set contains 20,234 cells that passed QC with a total of 32,711 genes detected. In the original study, the cell types of the cells in the Pixel-seq data were annotated using the cell-type annotation of a scRNA-seq data set^35^ as reference. We used these cell-type annotations in our analyses. For the Pixel-seq data, the raw counts were normalized by the total number of transcripts using the “normalize_total” function in Scanpy, and then log-normalized with “log1p” in Scanpy.

We downloaded mouse olfactory bulb scRNA-seq data^35^ from Gene Expression Omnibus (accession number: GSE121891), based on which the Pixel-seq data was annotated in the original study^29^. The processed scRNA-seq data contains 51,426 cells that passed QC and annotated in the original study^35^ with a total of 18,560 genes detected. We used cell-type annotations provided by the original study in our analyses. To simplify the original annotation, cell subtypes were merged consistently for both the Pixel-seq data and this scRNA-seq data. The scRNA-seq data were normalized in the same way as the Pixel-seq data.

### Analyses for demonstrating the transcript leakage problem

#### Exploratory data analysis

As we observed gene expression variations across brain regions, we further focused our analysis to the hippocampus formation (HPF) region, which has a clean structure of Cornu Ammonis (CA) and Dentate Gyrus (DG) primarily composed of neurons, for certain analyses. Specifically, the analyses shown in Figure 1C-E,G-L were all based on data from the HPF region only. The other analyses, shown in Figure 1A-B,F,M-O, and Figure 5, were obtained from the whole brain instead of only the HPF region.

For the other three real ST data sets, we used the whole tissue section to demonstrate the transcript leakage problem.

#### Definition of exclusively expressed genes

We identified a list of marker genes exclusively expressed in each major cell type using the companion scRNA-seq data from the HPF region as reference for the mouse brain MERFISH data set. The exclusively expressed genes for each cell type were defined as genes whose transcripts were detected in at least 80% of this cell type and at most 10% in each of the other cell types. We compared their gene expression in the MERFISH and scRNA-seq data. We further formed mutually exclusive gene pairs (*g1, g2*) whereas *g1* and *g2* are exclusively expressed marker genes of different cell types. We checked the co-detection of mutually exclusive gene pairs in the same cells as implication of contamination. To demonstrate the generality of the issue, we extended the analysis to cover 12 major brain regions (Olfactory areas, Cortical subplate, Hippocampus formation, Isocortex, Striatum, Pallidum, Hypothalamus, Thalamus, Midbrain, Pons, Medulla, and Cerebellum). A set of exclusively expressed markers for each major cell type across the brain region was identified firstly based on the scRNA-seq data, and then we prioritized for canonical markers and genes identified as exclusively expressed markers across multiple brain regions.

#### Analyses of transcript distance to cell centroid and cell distance to potential contamination source

We performed two distance-based analyses using the exclusively expressed marker genes. Both analyses assume that the expression of these markers in the corresponding cell type is endogenous expression, while the detection of them in other cell types is potential contamination. The first analysis aimed to check the relative closeness of endogenously expressed transcripts to cell centers compared to contamination transcripts. The centroid of each cell was estimated as the average coordinate of all transcripts assigned to this cell. The Euclidean distance between each transcript and its corresponding cell centroid was calculated. For each cell, the mean transcript-to-centroid distance was used as a normalization factor. Specifically, each transcript’s distance was divided by the cell-specific mean distance, such that a normalized value <1 indicates the transcript is located closer to the cell centroid than average, whereas a value >1 indicates it is farther from the centroid than average. Normalization is necessary to combine results across cells. The Welch’s t-test with multiple hypothesis testing correction adjusted by Bonferroni Correction was performed to check whether endogenous transcripts are significantly closer to cell centroid than potential contamination. The second analysis stratified the observed contamination transcript levels by distances to the contamination sources or cells endogenously expressing these transcripts. The basic assumption is that, if the contamination is due to transcript leakage, it should exhibit a diffusion-like spatial pattern. For cell type ***A*** and its exclusively marker gene ***g***, we correlated the detected level of ***g*** in cells of different cell types (as an approximation of contamination level) and their distances to the nearest cell of cell type ***A*** (as distance to potential contamination source).

#### Alternative segmentation strategies

We used three alternative approaches to perform cell segmentation in addition to the original cell segmentation for the mouse brain MERFISH data set. In the first approach, staring with the original segmentation, we drew the smallest circle centered at the centroid of the cell segment that covers all transcripts assigned to each cell, and then shrank its radius by 50%. In the second approach, we performed nuclear segmentation using poly-T staining signals only, as opposed to the original cell segmentation that considers both poly-T and DAPI signals. In both approaches, transcripts that still fall within the resulting segment of a cell are more likely to be genuinely expressed by that cell. In the third approach, we used another cell segmentation method, Baysor^36^, as opposed to the original CellPose method^37^.

Specifically, the original cell boundaries were predicted by the CellPose method as described in the paper^25^, using staining images of cellular components including DAPI staining of nuclei and PolyT staining of poly-adenylated RNA to approximate cytoplasm. As contamination due to transcript leakage exhibits a diffusion-like spatial pattern, transcripts closer to the cell centroids are more likely to be endogenous expressed than transcripts closer to the cell boundaries, according to our previous analysis. Therefore, the first method utilized the existing cell segmentation and shrunk the cell radius by half. We defined the cell radius as the longest distance from any transcript assigned to this cell to the cell centroid estimated by the segmentation method. Then, we excluded transcripts originally assigned to each cell but having a distance to the cell centroid more than half of the original cell radius. The remaining transcripts were counted to construct a new cell x gene expression matrix. Only two-dimensional coordinates (x, y) were used in this analysis as the CellPose method estimated the cell center in two dimensions excluding the Z-axis.

The second method is nuclei segmentation instead of whole cell segmentation. We called the CellPose method through the VizGen postprocessing tools (VPT v1.0.2) to segmentation cell nuclei based on DAPI staining images from all 7 Z-stacks. The segmented cells were mapped to annotated cells reported in the original study to infer their cell types.

The third method is by segmenting cells using a transcript distribution-based method called Baysor. The Baysor (v0.6.2) segmentation took locations of transcripts from all 7 Z-stacks as input. The minimum number of molecules/transcripts per gene and per cell were set to 50 and 20 respectively to run the Baysor method. The cell radius was estimated as sqrt(area/pi) where the area is the “area of the convex hull around the cell molecules” output by Baysor. Cells with an estimated radius of less than 3 microns and larger than 10 microns were filtered out. The remaining cells were mapped to annotated cells reported in the original study to infer their cell types.

To infer cell-type annotation of the new segmentation and enable direct comparison between two segmentations, we aligned the new segmentation (nuclei-only CellPose segmentation and Baysor segmentation) to annotated cells reported in the original study segmented by CellPose for the whole cell body. For each cell in the new segmentation, we found the nearest annotated cell within a certain distance between their cell centers (2 microns for nuclei-only CellPose segmentation and 10 microns for Baysor segmentation). A successful alignment requires that at least 80% of the cell area as defined by the bounding box is covered by the other cell. We excluded mappings when more than one cell mapped to the same annotated cell. For remaining mappings, we transfer the cell-type annotation from the annotated cell to the newly segmented cell. For cells in the HPF region of a MERFISH brain tissue sample (C57BL6J-638850.38), 8,029/10,160 (79.0%) annotated cells matched with cell nucleus segmented by CellPose. 3,331/10,160 (32.8%) annotated cells matched with cells by Baysor segmentation.

The data normalization follows the same practice as in the original study. The raw count gene expression matrix was normalized by cell volume for CellPose segmentation and cell area by Baysor segmentation. The normalized values were scaled up by multiplying 1,000 and then taken log2 with the pseudo count being 1.

### Details of the DeLeakage method

#### Summary of the DeLeakage model

The hierarchical model used by DeLeakage to estimate gene expression and diffusion parameters is as follows:

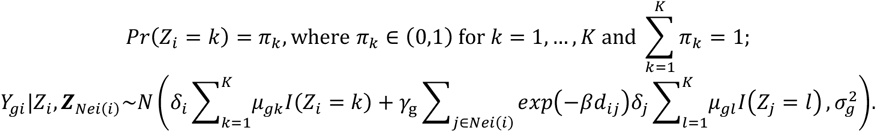

Collectively, ***Y*** = (***Y***_1_, …, ***Y***_*N*_), where ***Y***_*i*_ = (*Y*_1*i*_, …, *Y*_*Gi*_ )^*T*^, are the observed data; ***Z*** = (*Z*_1_, …, *Z*_*N*_) are the missing data; and the parameters ***Θ*** = {π, (***μ***_1_, …, ***μ***_*K*_), ***γ***, *β*, ***∑***} consist of the cell-type proportions π = (*π*_1_, …, *π*_*K*_), baseline mean expression levels for each cell type ***μ***_*k*_ = (*μ*_1*k*_, …, *μ*_*Gk*_ ), the gene-specific variance 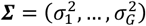, the gene-specific diffusion parameter ***γ*** = (*γ*_1_, …, *γ*_*G*_), and the decay parameter with respect to distance *β*.

#### Model identifiability

*Proof of Theorem 1*. Given *Pr*(***Y***|***Θ***) = *Pr*(***Y***_1_, …, ***Y***_*N*_|***Θ***), the marginal probability density function *f*(***Y***_*i*_|***Θ***) for the observed expression ***Y***_*i*_ for cell *i* is a mixture of high-dimensional normal distributions. Let *n*_*i*_ = |*Nei*(*i*)| be the number of neighbors of cell *i* and *Nei*(*i*)[*s*] be the *s*-th element of *Nei*(*i*). As *Z*_*i*_ has *K* possibilities and each *Z*_*j*_ for *j* ∈ *Nei*(*i*) has *K* possibilities, *f*(***Y***_*i*_|***Θ***) consists of a total of 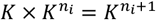 components. Let *A* = {1, …, *K*} and 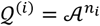, and let 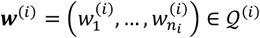 be a specific configuration of ***Z*** _*Nei* (*i*)_, where 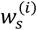 corresponds to *Z* _*Nei* (*i*) [*s*]_. Consequently, each component of *f*(***Y***_*i*_|***Θ***) corresponds to a configuration of (*Z*_*i*_, ***w***^(*i*)^) and is a multivariate normal distribution whose covariance matrix is 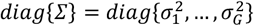 and whose mean for each dimension (i.e., each gene) is 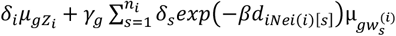. With an abuse of notation, we use *N*(*x*; *μ, τ*^2^) to represent the probability density function for the normal distribution *N*(*μ, τ*^2^) evaluated at *x*. As a result,

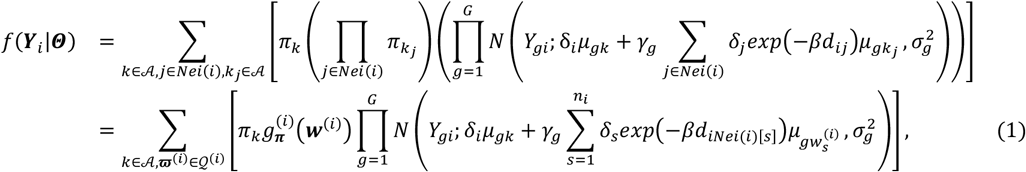

where 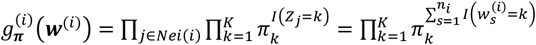.

Because the multivariate normal mixture model is identifiable up to label switching so that 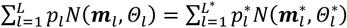 if and only if 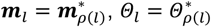, and 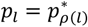, for *l* = 1, …, *L* ^38^, Equation (1) and *Pr*(***Y***|***Θ***) = *Pr*(***Y***|***Θ***^∗^) give that 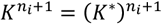 so that *K* = *K*^∗^ and there exists a permutation

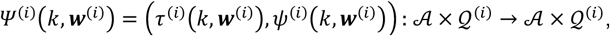

where *τ*^(*i*)^(*k*, ***w***^(*i*)^): *A* × *Q*^(*i*)^ → *A* and 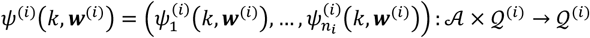, such that

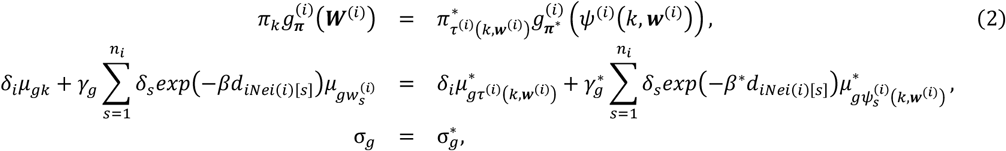

for *i* = 1, …, *N, g* = 1, …, *G, k* ∈ *A*, and ***w***^(*i*)^ ∈ *Q*^(*i*)^. For a given cell type *k*, the mean expression level of its marker gene *g*^(*k*)^ for the mixture component of the configuration 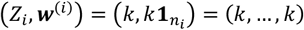 under ***Θ*** is:

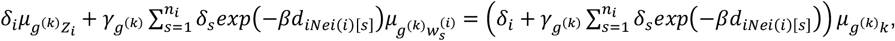

which is greater than 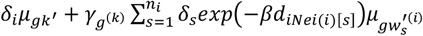 for any other configuration (*k*^′^, ***w***^′(*i*)^) according to Condition (C1). Similarly, denote the cell type in which gene *g*^(*k*)^ has the highest expression level under the alternative parameterization ***Θ***^∗^ as ρ(*k*), then under ***Θ***^∗^, the mixture component of *f*(***Y***_*i*_|***Θ***) that has highest mean for gene *g*^(*k*)^ must be 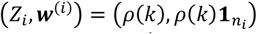 and 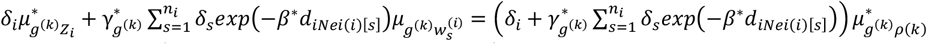, which equals 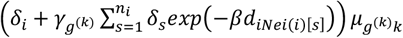. Consequently, 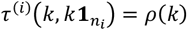 and 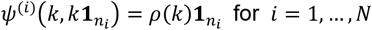. As a result, from Equation (2), we obtain 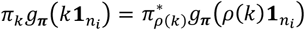 so that 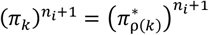 and hence 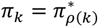. Meanwhile, for any gene *g*, we have

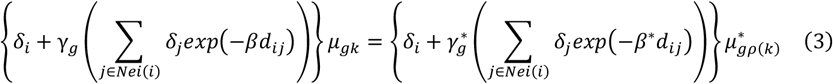

In particular, for the marker gene *g*^(*k*)^, Equation (3) leads to 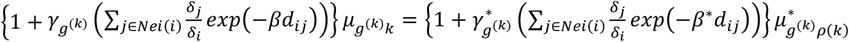 . Now by taking cells *i*_1_, *i*_2_ and *i*_3_ mentioned in Condition (C2), we obtain

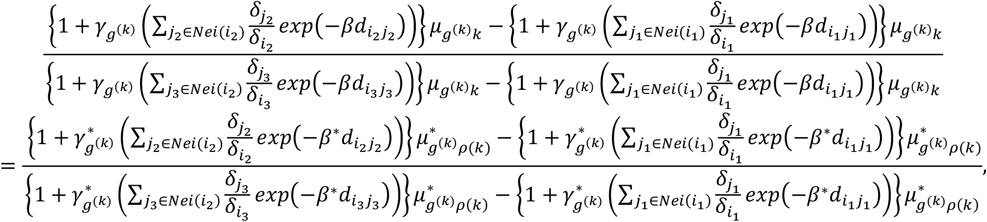

which result in 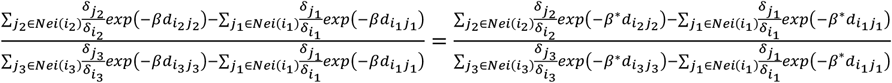, so that 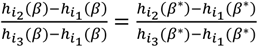 and hence *h*(β) =h(β^∗^). According to Condition (C2), h(x) is strictly monotone, and therefore, *β* = *β*^∗^ . Notice that Condition (C2) also implies that 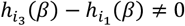. Consequently, for each gene *g*, by comparing the corresponding equations of Equation (3) for cells *i*_1_ and *i*_3_, we can obtain that

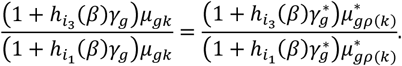

As a result, 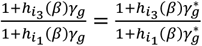 so that 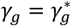 for each gene *g* . After plugging in 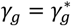 into Equation (3), we have ***μ***_*k*_ = ***μ***_*ρ*(*k*)_. Thus, the DeLeakage model is identifiable up to label switching.

##### An example situation in which Condition (C2) is fulfilled

For the extreme case when all the cells’ volumes are the same, say 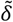, and the distances between a given cell *i* and all of its neighboring cells are identical, say *d*_i_, then 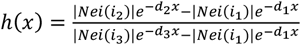 so that as long as the distances satisfies *d*_3_ ≥ *d*_1_ ≥ *d*_2_ or *d*_3_ ≤ *d*_1_ ≤ *d*_2_ and the number of neighboring cells are different, excluding the degenerate case where *d*_3_ = *d*_1_ = *d*_2_, Condition (C2) is fulfilled.

#### Prior and posterior distributions

We assign independent priors to each component of ***Θ*** and let 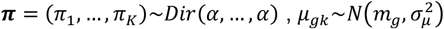 for *g* = 1, …, *G* and *k* = 1 …, *K, γ*_*g*_∼*B*e*ta*(*a*_*γ*_, *b*_*γ*_) for *g* = 1, …, *G*, and 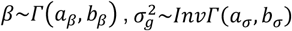 for *g* = 1, …, *G*. Consequently, the full posterior distribution becomes:

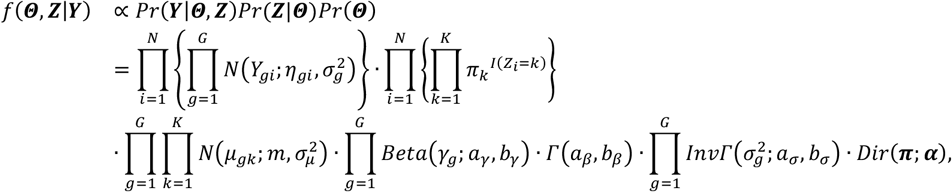

where 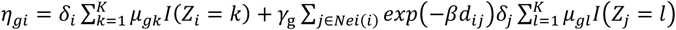.

### Posterior inference

To sample from the posterior distribution of each parameter, we developed an MCMC algorithm built upon the Gibbs sampler^39^. At any iteration *t* + 1:

1. We sample 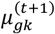 from 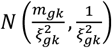, where:

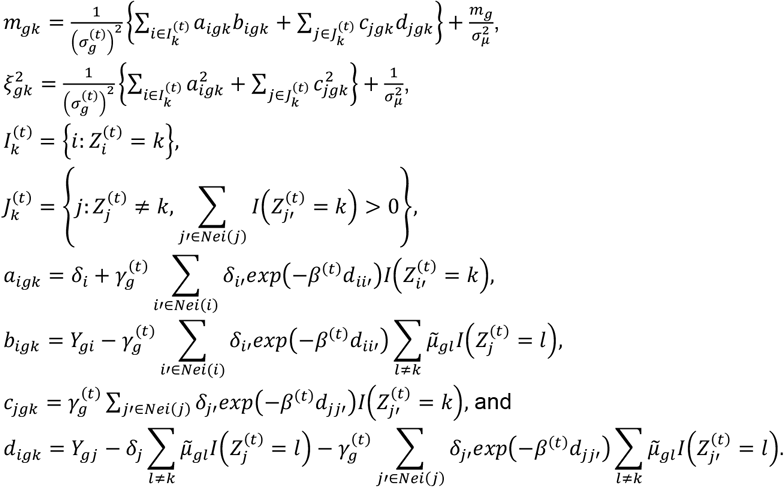

Due to the sequential sampling of *μ*_*gk*_ for *k* = 1, …, *K*, we denote 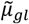 as the current state of *μ*_*gl*_ when updating *μ*_*gk*_ for brevity.
2. For the gene-specific contamination parameter, we update 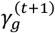 by a Metropolis-Hastings step with the proposal as 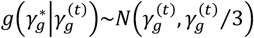 or 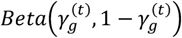 . The procedure of generating candidates is as follows:
  a. Generate 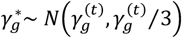 and keep it if the sampled value is between 0 and 1;
  b. Otherwise, generate 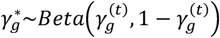. Denote 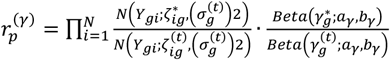, where 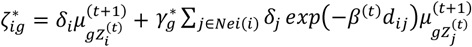 and 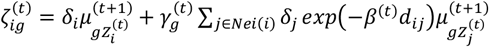. We accept 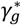 with probability *ρ* = *min*{*r*^(*γ*)^, 1}, where:
    - If 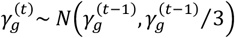 and 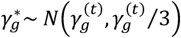, we have 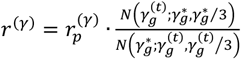;
    - If 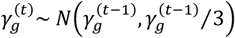 and 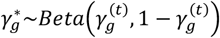, we have 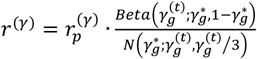;
    - If 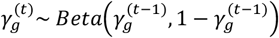 and 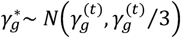, we have 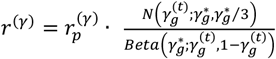;
    - If 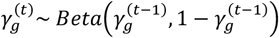 and 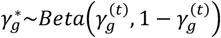, we have 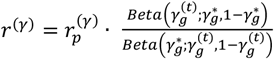.
3. For the distance decay contamination parameter, we update *β*^(*t*+1)^ by a Metropolis-Hastings step with the proposal as *g*(*β*^∗^|*β*^(*t*)^)∼N(*β*^(*t*)^, *β*^(*t*)^/3) or Γ(*β*^(*t*)^/3, 1/3) . The procedure of generating candidates is as follows:
  a. Generate *β*^∗^∼N(*β*^(*t*)^, *β*^(*t*)^/3) and keep it if the sampled value is between 0 and 1;
  b. Otherwise, generate *β*^∗^∼ Γ(*β*^(*t*)^/3, 1/3). Denote 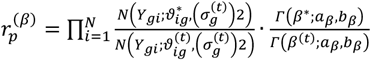, where 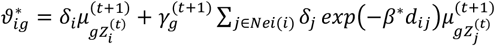 and 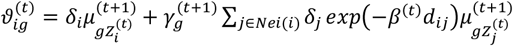. We accept β^∗^ with probability *ρ* = *m*i*n*{*r*^(*β*)^, 1}, where:
    - If *β*^(*t*)^∼ *N*(*β*^(*t*−1)^, *β*^(*t*−1)^/3) and *β*^∗^∼ *N*(*β*^(*t*)^, *β*^(*t*)^/3), we have 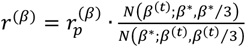;
    - If *β*^(*t*)^∼ *N*(*β*^(*t*−1)^, *β*^(*t*−1)^/3) and *β*^∗^∼ Γ(*β*^(*t*)^/3, 1/3), we have 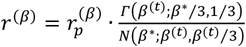;
    - If *β*^(*t*)^∼ Γ(*β*^(*t*−1)^/3, 1/3) and *β*^∗^∼ *N*(*β*^(*t*)^, *β*^(*t*)^/3), we have 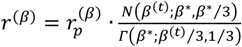;
    - If *β*^(*t*)^∼ Γ(*β*^(*t*−1)^/3, 1/3) and *β*^∗^∼ Γ(*β*^(*t*)^/3, 1/3), we have 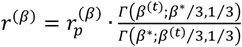;
4. We sample 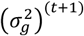 from 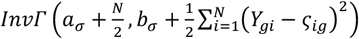, where 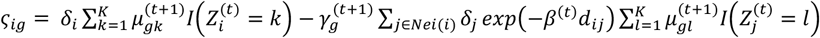.
5. For the cell type proportion, we sample π^(*t*+1)^ from Dirichlet distribution 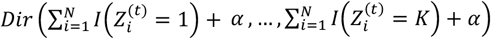.
6. For each cell *i*, we update its cell type label according to

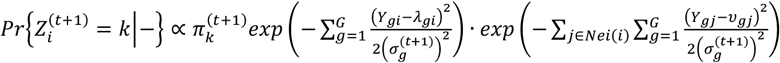

where 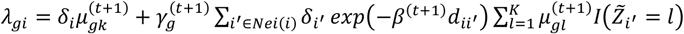 and 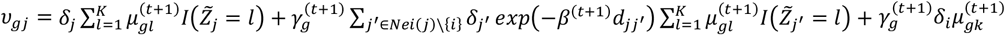 . Due to the sequential sampling of *Z*_i_ for *i* = 1, …, *N*, we denote 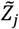 as the current state of *Z*_*j*_ when updating *Z*_*i*_ for brevity.

After the burn-in period, we can use the mode of the posterior samples of *Z*_*i*_ to infer the cell type for each cell and the mean of the posterior samples to estimate the other parameters. We compute corrected transcript level for gene *g* in cell *i* as 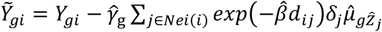 . If the computed value of 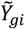 is smaller than zero, we set it to zero.

#### Parallel computation

The large data volume of ST data calls for parallel computing. Fortunately, unlike general MCMC algorithms constrained by sequential updates, steps (1), (2), and (4) can be computed in parallel for either all genes or all cells.

The parallelization for step (6), the update of cell-type label *Z*_*i*_, is more challenging because the full conditional distribution of *Z*_*i*_ depends not only on cell *i* ‘s gene expression profile but also on its neighbors’ gene expression profiles. Therefore, we propose to implement a scheme as shown in Figure 3. Suppose the maximum distance between any two neighboring cells is *r*. (a) First, we will randomly sample a cell from the entire tissue slice. After excluding this cell and all cells within its 2*r* neighborhood, we will sample a second cell. We will then exclude both the first and second cells, along with all cells within their 2*r* neighborhoods, and we will randomly sample a third cell. This process will be repeated until we run out of cells, and all the selected cells will be assigned to layer 1. (b) After excluding cells from layer 1, we will repeat the process to select cells for layer 2. (c) After excluding cells from layers 1 and 2, we will select cells for layers 3. We will repeat the process until we can select fewer than 100 cells; we will denote the resulting layer as layer *M*. Subsequently, we will collect all remaining cells to be grouped into layer *M* + 1. (d) As cells within the same layer for layers 1 to *M* will not be each other’s neighbor and hence not in each other’s Markov blankets, their *Z*_*i*_ values can be updated in parallel. (f) For cells in layer *M* + 1, we will update their *Z*_*i*_’s sequentially.

For GPU computation, we further leverage the reduction technique, which takes advantage of the architecture of GPU. For example, to sum over 1,204 numbers using reduction, in cycle 1, every two of the 1,024 numbers will be summed in parallel to produce 512 values; in cycle 2, the 512 values will be further added up and reduced to 256, and so on. Thus, the calculations in steps (3) and (5) can also be parallelized for summations over cells.

#### Hyper-parameter settings when applying DeLeakage to simulated and real data

For each cell *i*, we define its neighborhood, *Nei*(*i*), based on a fixed radius *r* : any cell *j* with Euclidean distance *d*_*ij*_ < *r* is denoted as a neighbor (*j* ∈ *Nei*(*i*)). We choose the fixed value of *r* to ensure that the median number of neighbors across cells is a prespecified value. In both our simulation and real-data analyses, we set *r* such that the median neighborhood size is 10. For the cell size factor *δ*_*i*_, if the volumes of cells are not available, *δ*_i_ can be computed alternatively as 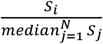, where 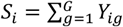 is the total read counts of cell *i* . The hyper-parameters of prior distributions are set as follow: *α* = 0.5,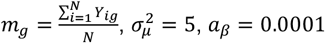, *b*_*β*_ = 1, *a*_*σ*_ = 1, *b*_*σ*_ = 1.

### Simulated data

An artificial tissue with 5 cell types was simulated, in which the different cell types roughly occupy distinct spatial territories but there are also cells that are present within the territories of other cell types (Figure 4A). Expression of 250 genes were simulated, which included i) genes with a high mean expression level in only one cell type, which mimic cell type markers, ii) genes with a high mean expression level in two cell types but at different levels, and iii) genes with the same mean expression level in all cell types, which mimic house-keeping genes (Figure 4B). For each cell, the detected transcript level of each gene was simulated by sampling a real expression level from the defined distribution followed by adding contamination caused by transcripts leaked by nearby cells according to the gene’s diffusion parameter.

Specifically, to generate the required patterns, we first uniformly sampled 20,000 points from a rectangle of size 2,400 × 2,000 and then kept the 15,261 points within the tissue region as the centroids of the cells. As a result, a total of *N* = 15,261 cells remained, and the median number of neighbors for each cell is 10. Next, after defining the five spatial domains, we randomly assigned cell-type labels to cells within each domain, with 80% probability coming from the primary cell type of each domain and 5% probability from each of the other four cell types. The cell size factor *δ*_*i*_ of each cell was sampled from *N*(1,0.01). The gene panel consists of *G* = 250, and the true underlying mean expression levels for each cell type are shown in Figure 4B. Specifically, the first 100 genes are set as the cell type marker genes that are expressed in only one cell type, and the numbers of marker genes for cell types 1-5 are set as 20, 25, 30, 10, and 15, respectively. The next 100 genes are set to be highly expressed in two cell types with differential expression between the two cell types, and these genes induce similarity between cell types. Finally, the last 50 genes represent housekeeping genes that are expressed in all cell types. The gene-specific variance 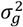 is set as 0.01 for all the genes. We sampled the gene-specific contamination parameters *γ*_*g*_ ‘s from a uniform distribution *U*(0.1,0.2) for *g* = 1, …,250 and set the remaining *γ*_*g*_ ‘s as 0.001 for *g* = 251, …,300 to mimic the set of genes that do not suffer from transcript leakage. The decay parameter of contamination *β* is set as 0.01 so that the range of *exp*(−*βd*_*ij*_) is between 0.7 and 1, according to what is observed from the real data. Under these three gene settings, we first simulated the underlying true expressions determined only by cell-type effects. We then generated the observed expressions by adding transcript leakage from neighboring cells. As a result, due to transcript leakage, while the spatial patterns of the true underlying expression levels of the marker genes can clearly show the five spatial domains (Figure 4F, Supplementary Figure 3A,F,K,P), the spatial patterns of the observed transcript levels of the marker genes are much more ambiguous (Figure 4G, Supplementary Figure 3B,G,L,Q).

### Comparisons with ST data denoising methods

#### The methods compared

We compared DeLeakage with two ST data denoising methods, SPLIT (v0.1.0) and SpotClean (v1.4.1). SPLIT was designed for *in situ* ST data at subcellular resolution. SpotClean was designed for removing noises in spot-based ST data with background spots which are supposed to have no expression.

#### Applying SPLIT

The SPLIT method^23^ assumes that noisy signals from neighboring cells of different cell types can be removed by estimating the proportion of different cell types forming the observed gene expression through cell type decomposition. It relies on a third-party cell-type decomposition method, RCTD^21^, to identify the primary and secondary cell types forming each ST cell, based on a scRNA-seq reference data set. SPLIT ran RCTD decomposition in the “doublet” mode. SPLIT further postprocessed the RCTD decomposition results and further inferred the corrected counts using the “purify” function with “DO_purify_singlets=TRUE”, which scales the observed expression to a purified expression level with the scaling factor being the primary cell type’s expected fraction of transcripts.

For the simulation study, the reference single-cell RNA-seq data were simulated to include all five cell types. For each cell type, 600 cells were generated, resulting in a total of 3,000 reference cells. The gene expression values for these reference cells were generated using the same cell-type mean expression levels and gene-specific variance parameters as those used to simulate the underlying true gene expression in the simulated spatial transcriptomics data, while there is no contamination or leakage effect added to the reference single-cell data. Therefore, the simulated reference data represent clean cell-type-specific expression profiles. Then, SPLIT adjusts the observed expression levels through a deconvolution-based procedure.

For the adult mouse brain MERFISH data^25^, the scRNA-seq data, generated in the same study using 10x Genomics Chromium Single Cell 3’ v3 protocol, was used as the scRNA-seq reference data which is required to run RCTD. The original data contains around 2.3 million cells, which imposes a big challenge as running RCTD on all scRNA-seq data uses more than 1TB RAM. To constrain the resources usage to a practical level, the scRNA-seq data was randomly down-sampled to 234K cells, around 1/10 of its original cell number. After down-sampling, each major cell type still has over 8K cells. The RCTD was run in the major cell type level. The resulting RCTD estimates, including cell-type proportions and cell-type-specific mean expression levels, were subsequently used by SPLIT to deconvolute and adjust the observed expression levels.

#### Applying SpotClean

SpotClean^20^ aims to tackle spot swapping, which happens when messenger RNAs bleed between nearby spots and hence cause contamination of expression levels, for spot-based ST methods. SpotClean assumes a transcript diffusion model with a global swapping rate and leverages background spots that are supposed to have no expression to estimate the model parameters and remove contamination. We applied SpotClean to image-based ST data as follows. Firstly, SpotClean requires empty or background spots to estimate background signals. To simulate background spots for image-based ST data, we used transcripts not assigned to any cells as background transcripts, and sampled spots of sizes similar as cells in the tissue slide, and we aggregated background transcripts to generate the gene expression profile for each spot as background spot. Following the suggestion by the original authors, we set the background spot ratio to 25% of all cells or spots. Secondly, SpotClean assumes that all spots have the same sizes. Therefore, for each cell in image-based ST data, we normalized the raw read counts by cell volume or total transcripts and used the normalized gene expression as the input of the SpotClean method.

For the simulation study, the background spot expressions are generated as follows. First, we used the filtered-out locations (4,739 locations) to represent the background spots, which are outside of the tissue region. Second, each location was contaminated by transcripts leaked from nearby locations, consistent with the assumption adopted by SpotClean. Third, the expression at each background location was generated only based on contamination. Different from the generation of observed expressions within the tissue region, endogenous gene expression was omitted for background spots, because these locations were assumed not to contain cells or intrinsic biological signals. Using both the observed tissue-region expression and the simulated background expression as input, SpotClean then adjusted the observed expression levels.

For the adult mouse brain MERFISH data^25^, we simulated the background spots by partitioning the tissue slide into 10 µm x 10 µm x 10 µm voxels in three dimensions (X, Y and Z) accordingly. Because the thickness (Z-dimension) of the tissue slide is around 10 µm, the partition was in fact performed in two-dimensional space (X and Y). The volume of each voxel is 1000 µm^3^. We calculated number of transcripts falling in each voxel. We filtered out voxels potentially falling outside the tissue region, which were defined as these having zero transcript in itself, and also having zero transcript in any of its six surrounding voxels. To avoid background voxels overlapping too much with cell area, we further filtered out voxels with over half of transcripts falling within the voxel and being assigned to segmented cells. The remaining voxels were used as background voxels or spots. For each background voxel, the transcripts falling within the voxel and not being assigned to any cells were aggregated to generate its observed transcript count profile. The centroid of each voxel was used as its coordinates to run SpotClean. Furthermore, due to the extremely high RAM usage (closed to 1 TB RAM) of SpotClean on the whole tissue region, we selected a sub-region for the analysis.

#### Performance measures

We systematically evaluated the performance of DeLeakage and the two ST denoising methods in terms of signal specificity and effects on downstream analyses in both cell type-agnostic and cell type-aware manner.

Signal specificity was evaluated using exclusively expressed marker genes, the exclusiveness of which were validated in the reference sc/snRNA-seq data. Due to the transcript leakage, the exclusively expressed marker genes are observed in other unrelated cell types. To evaluate signal specificity, we used Jaccard index of exclusively expressed gene pairs where each gene pair (g1, g2) are expected to be exclusively expressed marker genes of two different cell types. Calculation of Jaccard index does not need cell-type annotation of ST data, and so is cell-type agnostic. The value of Jaccard index ranges from 0 to 1, with 0 indicating perfect exclusiveness and 1 indicating co-observing in every cell.

The effect of denoising on cell-type annotation was evaluated by calculating Cosine similarity between each pair of cell types using their average gene expression profiles. Lower cosine similarity indicates greater transcriptional divergence between cell types, suggesting improved separability for downstream cell-type annotation.

#### Performance measures specifically for simulation data

For the simulation study, the underlying true expression is available, allowing us to calculate the residuals between the decontaminated expression and the true expression. Here we denote *Ŷ*_*gi*_ as the adjusted expressions and then the residuals of expressions are defined as *r*_*gi*_ = *Y*_*gi*_ − *Ŷ*_*gi*_. For each gene *g*, we can have different distribution of *r*_*gi*_ from different methods and then used the boxplots and violin plots to display. Residual distributions more concentrated around zero indicate that the decontaminated expression is closer to the underlying true expression.

Furthermore, based on the predefined gene-panel design, including marker genes and DE genes, we conducted pairwise t-tests between cell types. For example, marker genes are expected to be differentially expressed between their corresponding cell type and the other cell types, while showing no differential expression between unrelated cell-type pairs. A similar evaluation was performed for the predefined DE genes. For each cell type pair (*k*_1_, *k*_2_), we conduct the t-test for each gene, then applied the Benjamini–Hochberg (BH) procedure to adjust for multiple testing across all pairwise cell-type comparisons and evaluated the resulting false discovery rate (FDR) under two different thresholds, 0.01 and 0.05. A lower FDR indicates better performance, as it reflects fewer spurious differential-expression signals after decontamination.

### Downstream analysis tasks

#### Post-clustering and ARI analysis

To perform cell-type annotation, for each data set, we applied a standard algorithm to form *K* cell clusters, where *K* is the number of cell types in the cell-type annotation. We checked the consistency of these clusters with the authors’ original annotations using Adjusted Rand Index (ARI), where a higher value indicates better consistency. We further compared the ARI values with or without applying DeLeakage.

Specifically, we first normalized the raw observed data or the adjusted expression values produced by DeLeakage and SPLIT. The normalized values were then transformed to the log scale using log(*x* + 1), and K-means clustering was performed using *kmeans()*, a basic clustering function in R. For a fair comparison, the number of clusters was set to be the same as the number of majority cell types in the ground-truth annotations. This strategy provides a simple and consistent framework for evaluating the quality of adjusted expressions from different methods. Finally, we calculated the ARI between the K-means clustering labels and the ground-truth annotations, and used the ARI values to compare the performance of different methods.

#### Moran’s I spatial auto-correlation analysis

To study spatially dependent expression, we analyzed spatial auto-correlation of detected transcript levels in cells that the genes are not expected to express. If the expression level of a gene in a cell correlates with its expression in the nearby cells, auto-correlation is high (i.e., closer to the maximum Moran’s I value of 1). For example, if a gene is specifically expressed in cell type X, and cells of this cell type tend to cluster spatially, then auto-correlation of this gene across all cells would be high. On the other hand, if the cells of cell type X are excluded from the analyses, any detected expression of the gene in the remaining cells is just noise if there is no transcript leakage, and thus auto-correlation should be close to zero. Yet due to transcript leakage, if these cells are contaminated by transcripts of this gene leaked by cells of cell type X, auto-correlation can be high again. Therefore, we expect that in the raw data affected by transcript leakage, auto-correlation of genes not expected to be expressed is high, but with proper adjustment of the data, auto-correlation of these genes will be reduced.

Specifically, to determine the ability of DeLeakage in reducing contamination levels of genes across different cell types and data sets, we performed spatial auto-correlation analysis using exclusive marker genes specific to each cell type. We compared raw count and decontaminated data from both our method and SPLIT for MERFISH mouse brain, MERFISH human heart and Xenium mouse brain data sets. For each annotated cell type per data set, we extracted the subset of cells assigned to that type from the expression matrix and their spatial coordinates. The marker-gene expression matrix for that cell subset was transposed into a cells-by-genes AnnData object with spatial coordinates stored. We constructed a spatial neighbor graph using scanpy.pp.neighbors(n_neighbors=11) (Scanpy version 1.11.5), which identifies the 10 nearest Euclidean-distance neighbors per cell. Moran’s I was then computed for each marker gene using scanpy.metrics.morans_i(adata), which operates on the connectivities graph.

### Running time and RAM usage

For the comparison of runtime and memory usage, we used Intel Xeon Gold 6246R processors at 3.4 GHz and fixed the number of CPU cores at 24 for both SPLIT and the CPU version of DeLeakage. For the GPU version of DeLeakage, we used an AMD Rome 7742 CPU at 3.4 GHz and an NVIDIA A100 GPU. For both simulated and real data sets, the total number of iterations was set to twice the burn-in period to facilitate convergence. Specifically, we used 1,000 iterations for the simulated data sets and 10,000 iterations for the real data sets. We implemented real-time monitoring of wall-clock runtime and peak memory usage during program execution. Resource metrics, including system memory, GPU memory, and process resident memory, were recorded every 10 seconds. At the end of each run, a final summary was generated, reporting the total runtime, peak memory usage, and program exit status.

## Code availability

Details on reproducing our analysis and analyzing other data sets are available on GitHub at https://yip-lab.github.io/DeLeakage/. DeLeakage is available as a Python package at https://github.com/Yip-Lab/DeLeakage. Code to reproduce the analysis and figures are available at https://github.com/Yip-Lab/DeLeakage and https://github.com/Yip-Lab/DeLeakage-analysis.

## Acknowledgements

CHS, PDA, BR, QZ, and KYY are supported by National Institute on Aging of the National Institutes of Health under Award Number U54AG079758. KYY is additionally supported by National Cancer Institute of the National Institutes of Health under Award Numbers P30CA030199 and R01CA287114, National Institute on Aging of the National Institutes of Health under Award Number R01AG085498, National Institute of General Medical Sciences of the National Institutes of Health under Award Number R21GM159319, National Heart, Lung, and Blood Institute of the National Institutes of Health under Award Number R21HL177724, The V Foundation under Award Number V2025-028, and internal grants of Sanford Burnham Prebys Medical Discovery Institute. MS and CC are supported by National Cancer Institute of the National Institutes of Health under Award Number UH3CA275669. The content is solely the responsibility of the authors and does not necessarily represent the official views of the funding agencies.

## Author contributions

CHS and KYY conceived the study. CHS, YZ, SH-CC, LL, YW, and KYY performed data analysis. CHS, YZ, LL, YW, and KYY were involved in the original design of the DeLeakage, while YZ, LL, and YW derived the model details, conducted the statistical inference, developed the parallel computing scheme, and implemented the algorithms. YZ and YW proved the model identifiability. CMC, MGT, JF, CK, PDA, BR, MJS, and QZ provided ST data. CHS, SH-CC, CMC, MGT, JF, CK, PDA, BR, MJS, QZ, YW, and KYY produced biological interpretations. CHS, YZ, YW, and KYY wrote the manuscript. All authors read and approved the final version of the manuscript.

## Supplementary Materials

### Supplementary Results

#### Prevalence of transcript leakage across spatial transcriptomics data produced from different tissues, species, and experimental platforms

To study the generality of the transcript leakage problem, we analyzed several additional ST data sets, including 1) MERFISH data produced from the human heart^28^, 2) Xenium data produced from the mouse brain (demo data from 10x Genomics) and 3) Pixel-seq data produced from the mouse brain^29^. Together, these data sets cover data from two tissue types of two species, produced using three ST platforms including both imaging-based and sequencing-based methods. In the first two studies, scRNA-seq or single-nucleus RNA-sequencing (snRNA-seq) data were also produced from related samples. For the Pixel-seq data set, we used published scRNA-seq data of the mouse brain^25^ in our analyses. Since raw data were not available in all cases, we focused on the analyses that require only processed data.

In all three studies, the original cell-type annotations of the cells in the ST data were produced by combining ST and scRNA-seq data (Supplementary Figure 2A-C). Based on these annotations, in all three cases, unexpected transcripts are detected in the ST data, namely transcripts of genes that are supposed to be markers of another cell type (Figure 2A-C). In contrast, these unexpected transcripts are detected at much lower levels in the corresponding scRNA-seq/snRNA-seq data (Figure 2D-F). Again, we confirmed the co-detection of markers of different cell types in individual cells in the ST data (Figure 2G-I). Using genes supposed to be expressed in mutually exclusive cells, we found that cells in the ST data have much more frequent co-detection than in the scRNA-seq/snRNA-seq data (Figure 2J-L). We also observed dependency of unexpected transcript level on the distance to the potential contamination source (Figure 2M-O).

Overall, these results show that transcript contamination is a prevalent problem in ST data.

#### Comparing DeLeakage, SPLIT, and SpotClean on simulated data

We assessed how spatial contamination may affect cell-type annotation by causing cells of different cell types to appear artificially more similar to each other. When similarity was computed using the actual gene expression levels, cells from the same cell type are generally more similar to each other than to cells from other cell types (Figure 4P). With transcript leakage in the simulated data, different cell types became more similar to each other (Figure 4Q). The true cell type differences were largely restored by the expression level adjustments performed by DeLeakage (Figure 4R). In contrast, SPLIT could only partially remove the artificial cell-cell similarities caused by transcript leakage (Figure 4S) while SpotClean over-corrected by reducing levels of genes that are actually expressed in multiple cell types.

We also performed differential expression analysis between pairs of cell types. In the simulated data, class i and class ii genes are truly differentially expressed between some cell type pairs, while class iii genes are not. When a gene is not really differentially expressed between two cell types but is identified to have differential expression, that constitutes a false positive call. When we applied differential expression analysis directly to the observed data, which is affected by transcript leakage, there was a very high level of false positives even after correcting for multiple hypothesis testing (Figure 4U-V). After applying DeLeakage to remove the contamination, the false positive rate was controlled to the target levels. In comparison, the false positive rate remained high after applying SPLIT or SpotClean (Figure 4U-V).

#### Application of DeLeakage to the other three real ST data sets

For the mouse brain Xenium and human heart data sets, DeLeakage was again effective as evidenced by the three types of analyses (Supplementary Figures 6-7). In both cases, DeLeakage achieved much better adjustments than SPLIT. For example, after applying DeLeakage to the mouse brain Xenium data, the Astro-Epen markers *Acsbg1* and *Aqp4* became mostly detected in Astro-Epen cells only (Supplementary Figure 7C), but after applying SPLIT, these transcripts were still detected in most cells in all cell types (Supplementary Figure 7D). For the mouse brain Pixel-seq data, we focused on the detected transcript levels of marker genes. We found that after applying either DeLeakage or SPLIT, detected transcript levels of marker genes of unrelated cell types were reduced, but DeLeakage was more successful in retaining the levels of marker genes of the relevant cell types (Supplementary Figure 8). For example, in granule cells, the level of its marker *Meg3* was almost unchanged by DeLeakage, but its level was substantially reduced by SPLIT (Supplementary Figure 8A).

#### Time and memory efficiency of DeLeakage

We evaluated the running time and peak RAM usage of DeLeakage. First, using simulated data with different simulation settings, we assessed the scalability of DeLeakage with increasing number of cells or genes. For both the CPU and GPU versions, both the running time and peak RAM usage scaled linearly with both cell number and gene number (Supplementary Figure 10A-D).

Next, we compared these computational requirements of DeLeakage and SPLIT when they were applied to the real data sets. In terms of running time, SPLIT required less time than the CPU version of DeLeakage, but the GPU version of DeLeakage required less time than SPLIT (Supplementary Figure 10E). For example, for the mouse brain MERFISH data set, which contains expression data of 500 genes in 4 million cells, the GPU version of DeLeakage only took slightly more than an hour to run.

In terms of peak memory usage, both the CPU and GPU versions of DeLeakage required substantially less than SPLIT (Supplementary Figure 10F).

## Supplementary Figure Legends

**Supplementary Figure 1.**
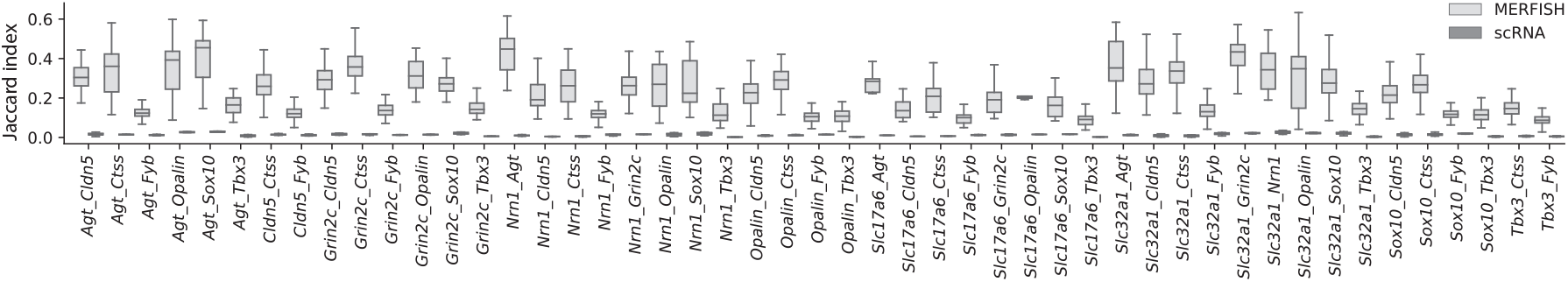
Co-detection of transcripts of gene pairs supposed to have mutually exclusive expression. Each gene pair is shown separately. For each gene pair, a Jaccard index is computed for cells in each brain region, and the box-and-whisker plot shows the distribution of Jaccard Index across all the brain regions based on either MERFISH data or scRNA-seq data.

**Supplementary Figure 2.**
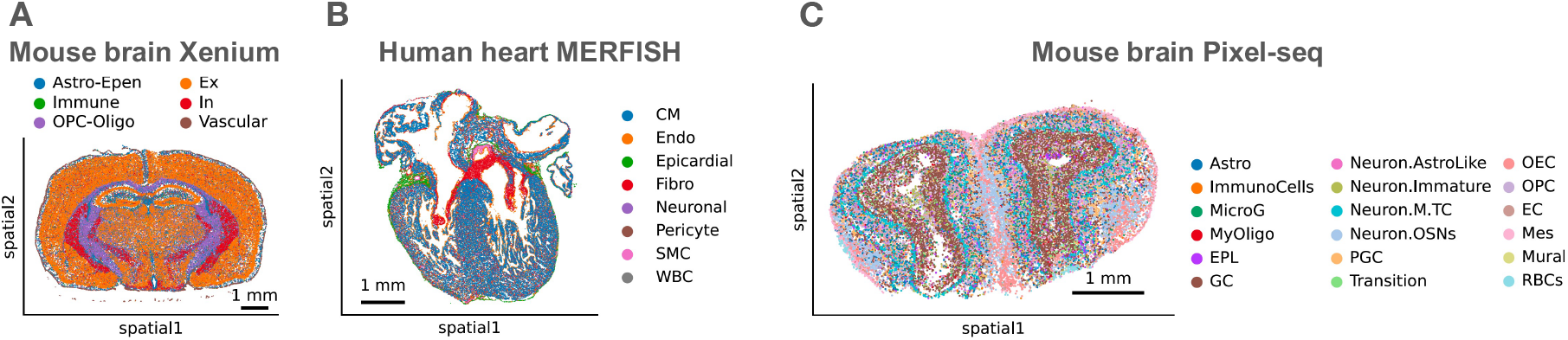
Spatial distribution of cell types in the additional spatial transcriptomics data sets. (**A**-**C**) Spatial distribution of cell types in human heart MERFISH data (A), mouse brain Xenium data (B), and mouse brain Pixel-seq data (C). Abbreviations of cell types in the mouse brain Xenium data set: Astro-Epen - astrocyte and ependymal cell; Ex - excitatory neuron; In - inhibitory neuron; OPC-Oligo - oligodendrocyte progenitor cell and oligodendrocyte. Abbreviations of cell types in the human heart MERFISH data set: CM - cardiomyocyte; Endo - endothelial cell; Fibro - fibroblast; SMC - smooth muscle cell; WBC - white blood cell. Abbreviations of cell types in the mouse brain Pixel-seq data set: Astro - astrocyte; ImmunoCells - immune cells; MicroG - microglia; MyOligo - myelinating oligodendrocyte; EPL - external plexiform layer interneuron; GC - granule cell; Neuron.AstroLike - astrocyte like neuron; Neuron.Immature - immature neuron; Neuron.M.TC - mitral and tufted neuron; Neuron.OSN - olfactory sensory neuron; PGC - periglomerular cell (a small inhibitory interneuron located in the glomerular layer of the olfactory bulb); Transition - transitional neuron; OEC - olfactory ensheathing cell; OPC - oligodendrocyte precursor; EC - endothelial cell; Mes - mesenchymal cell; Mural - mural cell; RBCs - red blood cells.

**Supplementary Figure 3.**
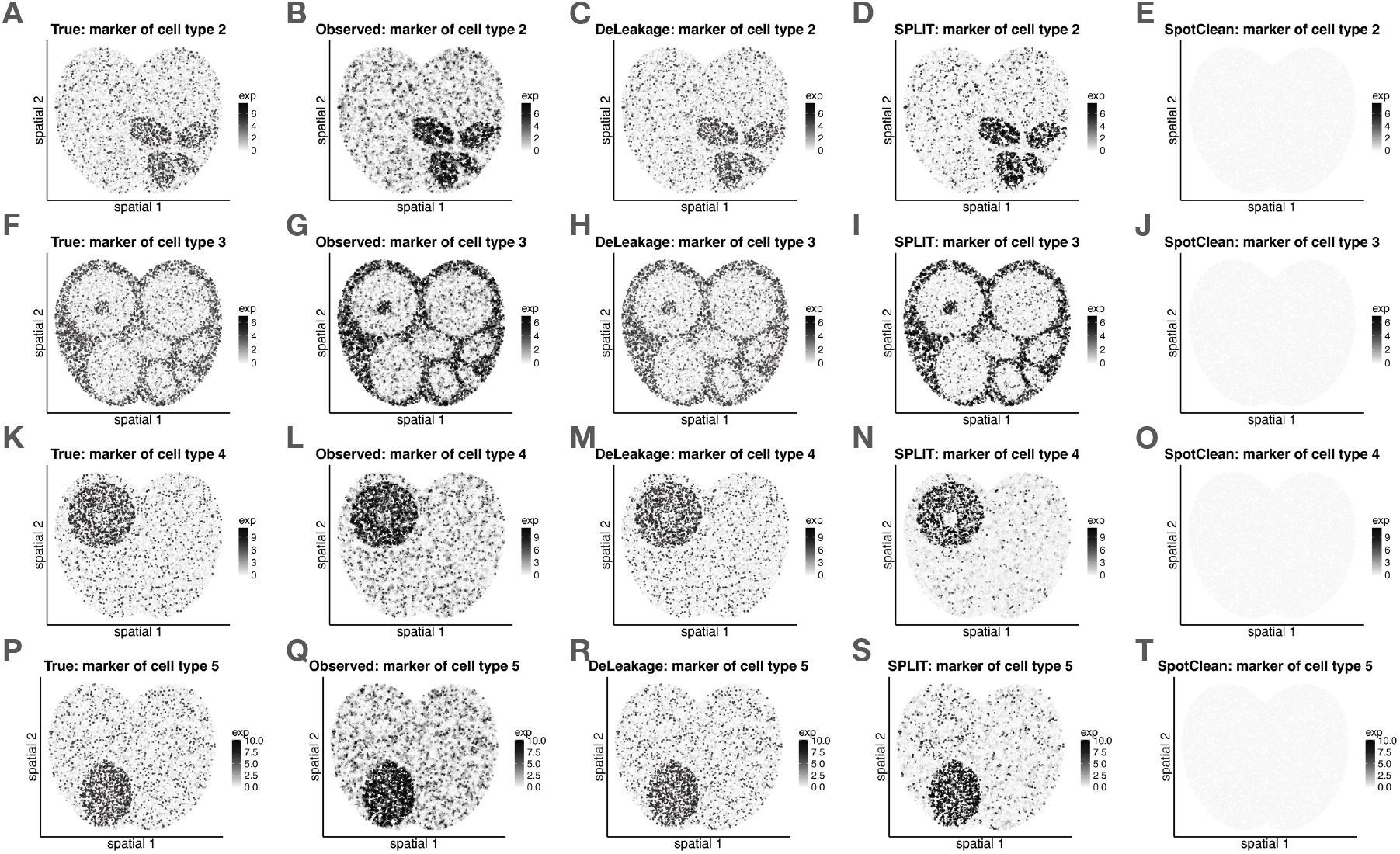
Benchmarking DeLeakage against other ST denoising methods based on simulated data (showing all cells). (**A**-**T**) Level of one of the marker genes in each cell type selected randomly in either the initial simulated data before transcript leakage, final simulated data after transcript leakage, the adjusted data after applying DeLeakage, SPLIT, or SpotClean. These results are shown in different columns, while each row shows the results for the marker genes of a different cell type.

**Supplementary Figure 4.**
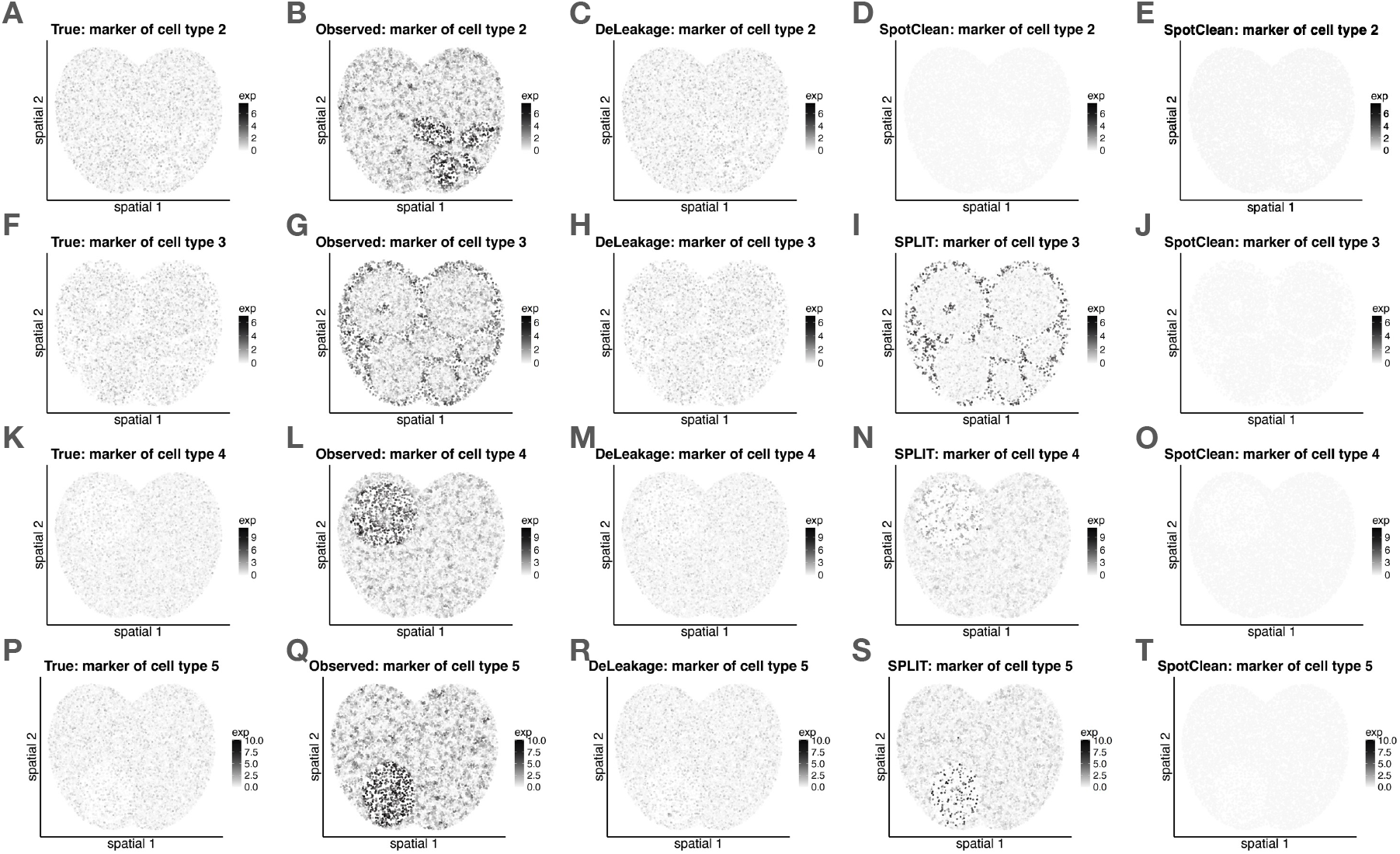
Benchmarking DeLeakage against other ST denoising methods based on simulated data (excluding cells of the cell type that expresses the marker gene). (**A**-**T**) Level of one of the marker genes in each cell type selected randomly in either the initial simulated data before transcript leakage, final simulated data after transcript leakage, the adjusted data after applying DeLeakage, SPLIT, or SpotClean. These results are shown in different columns, while each row shows the results for the marker genes of a different cell type.

**Supplementary Figure 5.**
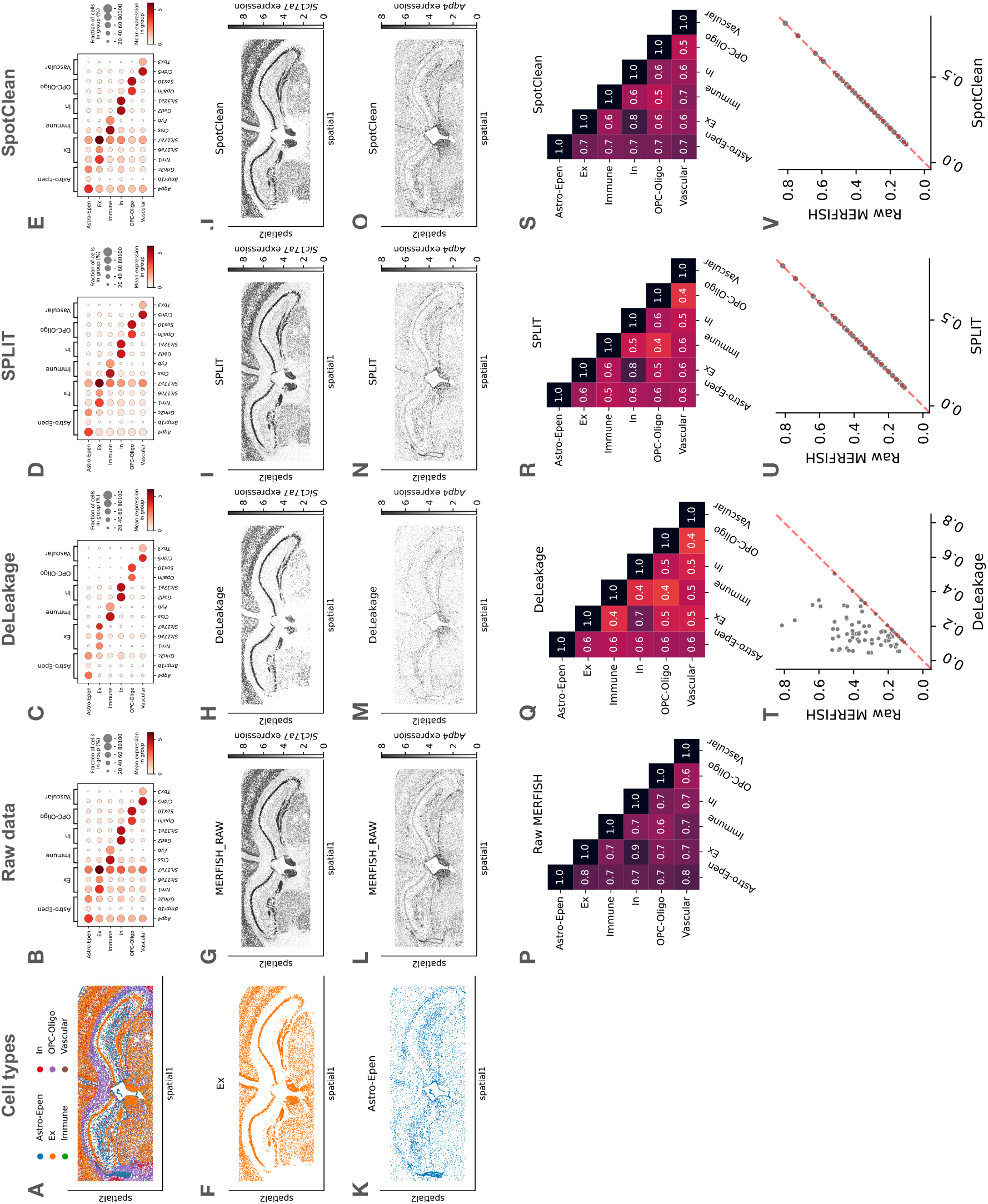
Benchmarking DeLeakage against other ST denoising methods based on a subregion in the mouse brain MERFISH data. (**A**) Spatial distribution of cell types. (**B**-**E**) Detection of transcripts of cell type-specific genes in the raw data (B), or after applying DeLeakage (C), SPLIT (D), or SpotClean (E). (**F**-**J**) Spatial distribution of excitatory neurons (F) and transcripts of its marker *Slc17a7* in the raw data (G), or after applying DeLeakage (H), SPLIT (I) or SpotClean (J). (**K**-**O**) Spatial distribution of Astro-Epen cells (K) and transcripts of its marker *Aqp4* in the raw data (L), or after applying DeLeakage (M), SPLIT (N) or SpotClean (O). (**P**-**S**) Cosine similarity of average gene expression among cell types, based on the raw MERFISH data (P), or after applying DeLeakage (Q), SPLIT (R), or SpotClean (S). (**T**-**V**) Co-detection of transcripts of gene pairs supposed to have mutually exclusive expression before and after applying DeLeakage (T), SPLIT (U), or SpotClean (V). For each gene pair, the Jaccard index is defined as the number of cells with both types of transcripts detected divided by the number of cells with either detected.

**Supplementary Figure 6.**
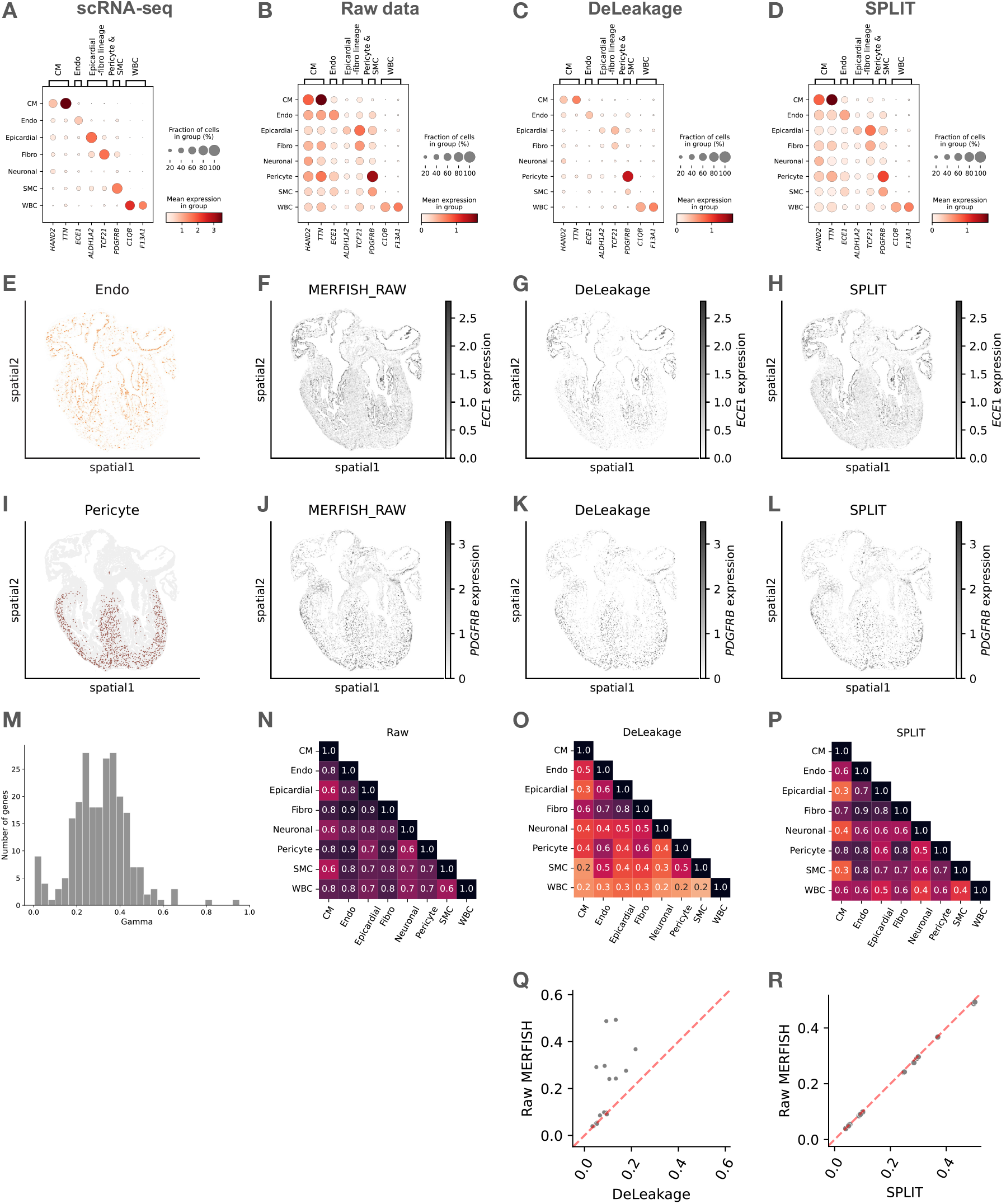
Improvement of transcript quantification in the human heart MERFISH data achieved by DeLeakage. (**A**-**D**) Detection of transcripts of cell type-specific genes in scRNA-seq data (A), raw MERFISH data (B), or after applying DeLeakage (C), or SPLIT (D). (**E-H**) Spatial distribution of endothelial cells (E) and transcripts of its marker *ECE1* in the raw data (F), or after applying DeLeakage (G) or SPLIT (H). (**I-L**) Spatial distribution of pericytes (I) and transcripts of its marker *PDGFRB* in the raw data (J), or after applying DeLeakage (K) or SPLIT (L). (**M**) Distribution of all genes’ diffusion parameters identified by DeLeakage. (**N**-**P**) Cosine similarity of average gene expression among cell types, based on the raw data (N), or after applying DeLeakage (O), or SPLIT (P). (**Q-R**) Co-detection of transcripts of gene pairs supposed to have mutually exclusive expression before and after applying DeLeakage (Q), or SPLIT (R). For each gene pair, the Jaccard index is defined as the number of cells with both types of transcripts detected divided by the number of cells with either detected.

**Supplementary Figure 7.**
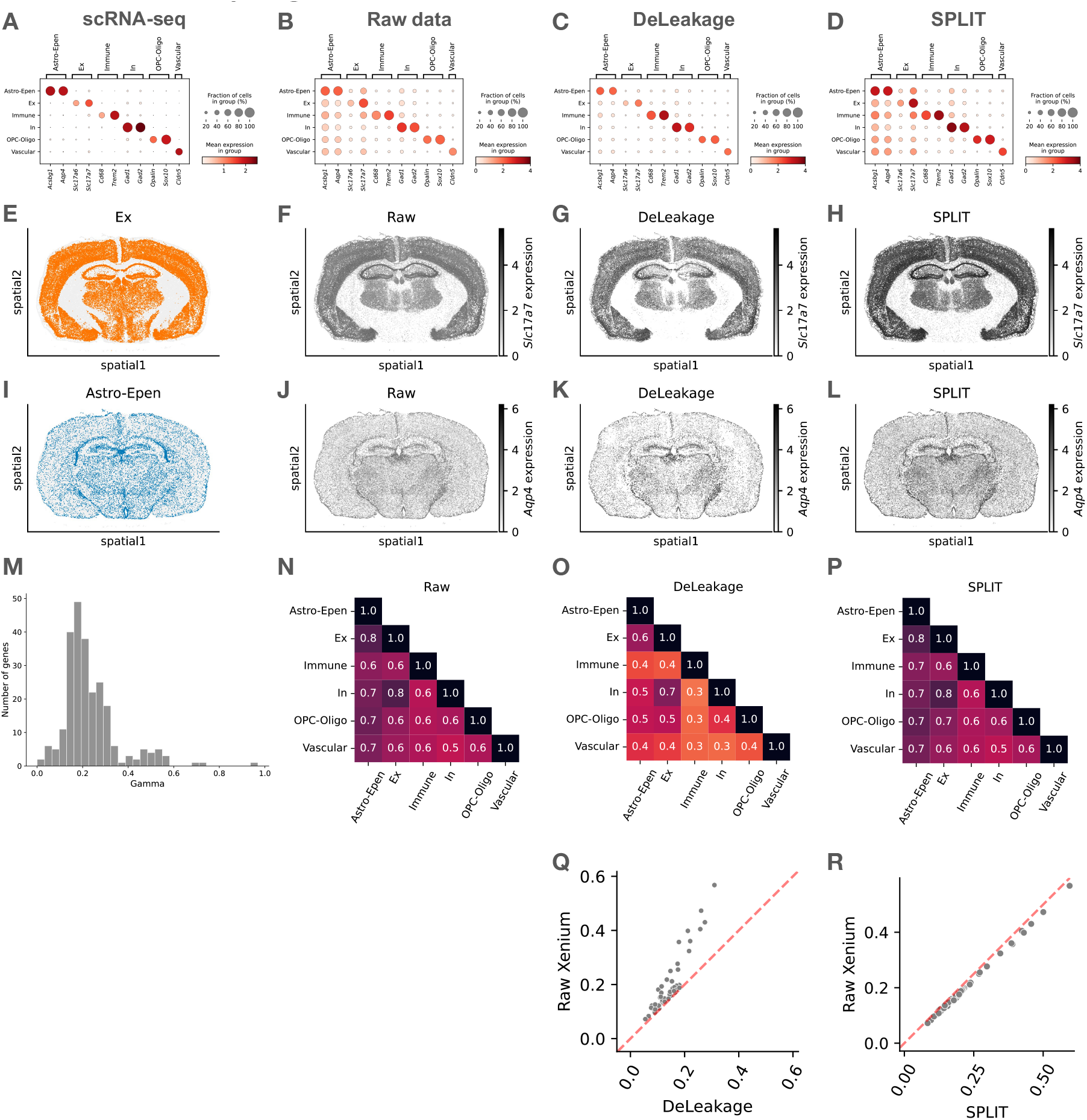
Improvement of transcript quantification in the mouse brain Xenium data achieved by DeLeakage. (**A**-**D**) Detection of transcripts of cell type-specific genes in scRNA-seq data (A), the raw data (B), or after applying DeLeakage (C), or SPLIT (D). (**E-H**) Spatial distribution of excitatory neurons (E) and transcripts of its marker *Slc17a7* in the raw data (F), or after applying DeLeakage (G) or SPLIT (H). (**I-L**) Spatial distribution of Astro-Epen cells (I) and transcripts of its marker *Aqp4* in the raw data (J), or after applying DeLeakage (K) or SPLIT (L). (**M**) Distribution of all genes’ diffusion parameters identified by DeLeakage. (**N**-**P**) Cosine similarity of average gene expression among cell types, based on the raw data (N), or after applying DeLeakage (O), or SPLIT (P). (**Q-R**) Co-detection of transcripts of gene pairs supposed to have mutually exclusive expression before and after applying DeLeakage (Q), or SPLIT (R). For each gene pair, the Jaccard index is defined as the number of cells with both types of transcripts detected divided by the number of cells with either detected.

**Supplementary Figure 8.**
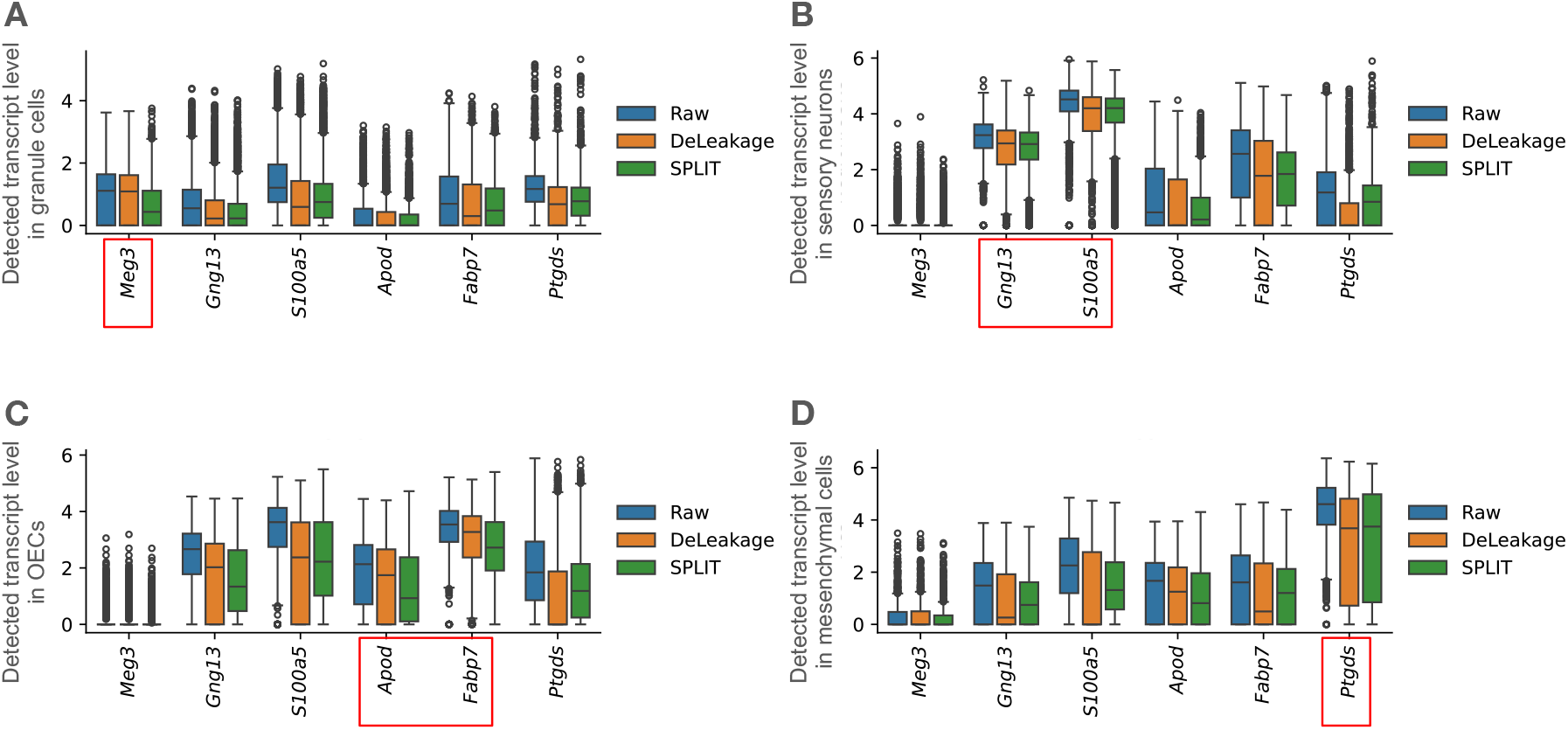
Improvement of transcript quantification in the mouse brain Pixel-seq data achieved by DeLeakage. (**A**-**D**) Detection of transcripts of genes in granule cells (A), sensory neurons (B), olfactory ensheathing cells (C), and mesenchymal cells (D) in the raw data or data adjusted by DeLeakage or SPLIT. For each cell type, the canonical markers of that cell type are highlighted in red boxes.

**Supplementary Figure 9.**
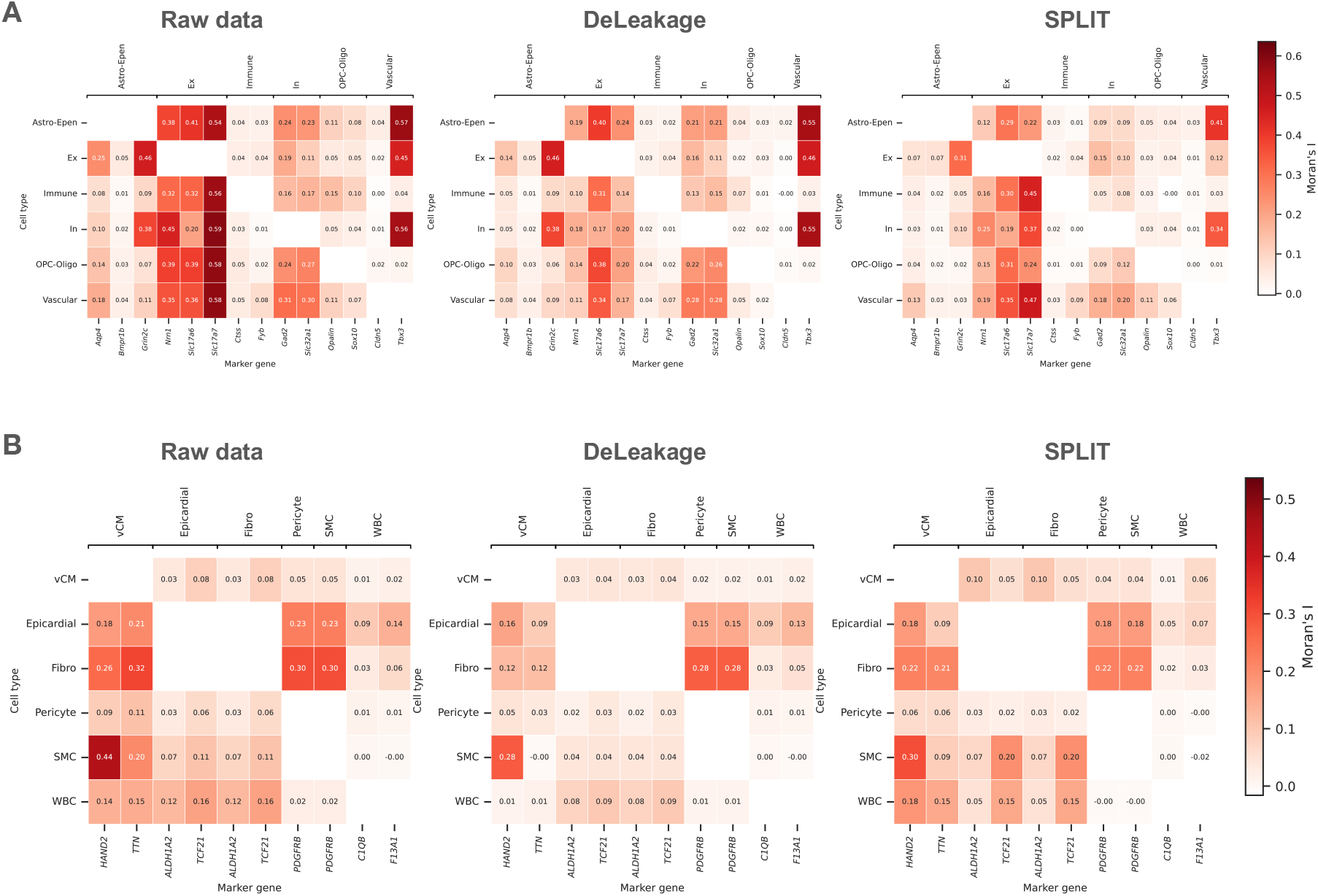
Improvement of spatially dependent expression calls achieved by DeLeakage. (**A**-**B**) Spatial auto-correlation of unexpected transcripts in the mouse brain MERFISH data (A) or human heart MERFISH data (B), based raw data (first column), DeLeakage-adjusted data (second column), or SPLIT-adjusted data (third column).

**Supplementary Figure 10.**
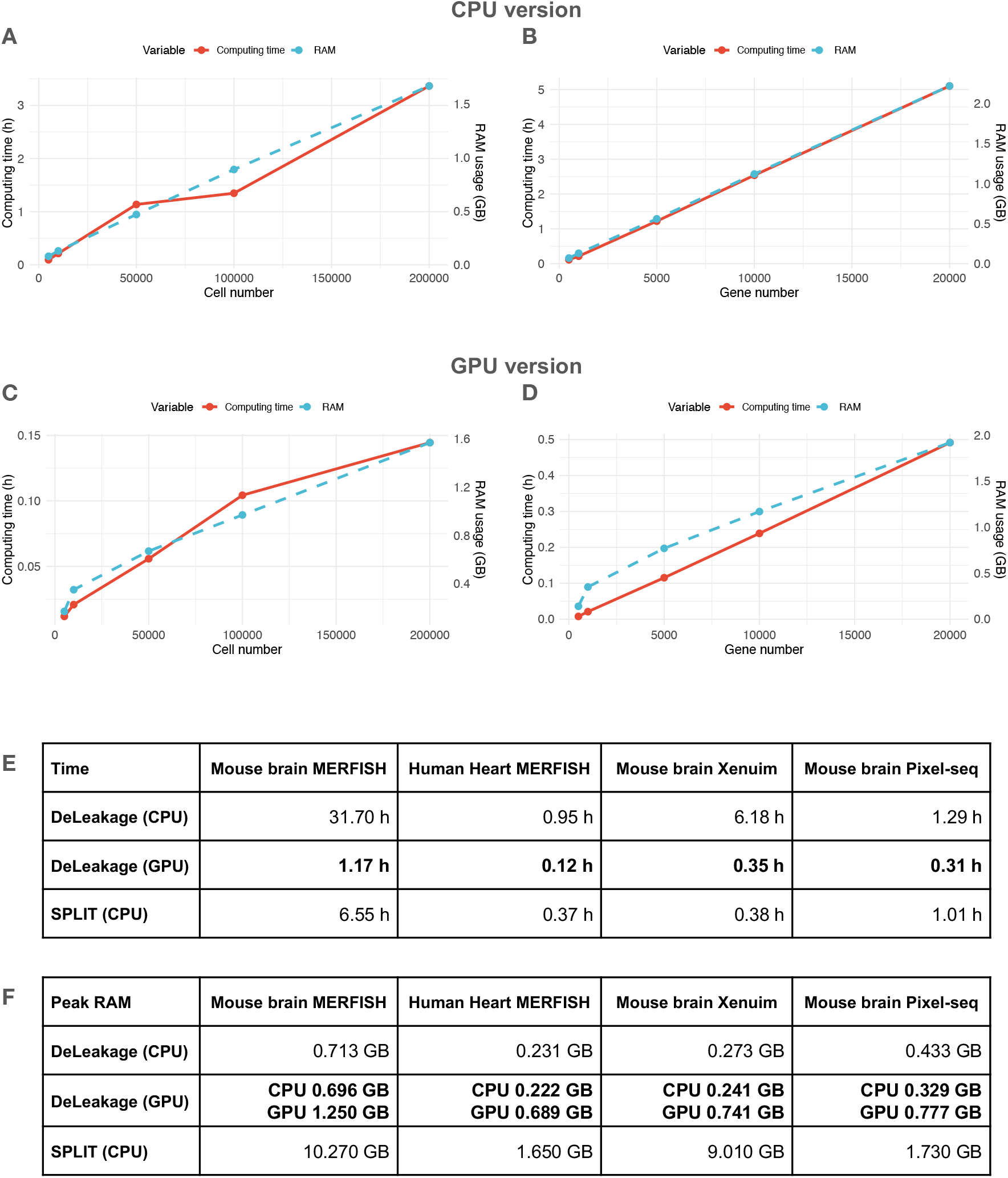
Running time and peak RAM usage. (**A**-**D**) Scalability of the CPU (A,B) and GPU (C,D) versions of DeLeakage with respect to increasing cell number and 1,000 genes (A,C) or increasing gene number and 10,000 cells (B,D). (**E**-**F**) Running time (E) and peak RAM usage (F) of the two versions of DeLeakage and SPLIT. For the GPU version of DeLeakage, both CPU peak RAM usage and GPU peak VRAM usage are reported.

